# 5’UTR translational inhibition of neuroblastoma dependency factors using the CR-1-31-B rocaglate

**DOI:** 10.1101/2024.08.29.610236

**Authors:** C Nunes, SL Bekaert, S Parsa, I Nelen, F Martens, E De Stanchina, E Sanders, E Hilgert, S T’sas, P Zhao, F De Vloed, A Eggermont, S Goossens, A Sablina, L Depestel, HG Wendel, K Durinck

**Affiliations:** Department for Biomolecular Medicine, Ghent University, Medical Research Building (MRB1), Corneel Heymanslaan 10, B-9000 Ghent, Belgium; Cancer Research Institute Ghent (CRIG), B-9000 Ghent, Belgium; Cancer Biology and Genetics, Memorial Sloan-Kettering Cancer Center, New York, NY, USA; Department of Diagnostic Sciences, Ghent University, Ghent, Belgium; VIB-KU Leuven Center for Cancer Biology, VIB, 3000 Leuven, Belgium; Department of Oncology, KU Leuven, Herestraat 49, 3000 Leuven, Belgium

**Keywords:** neuroblastoma, precision oncology, translation, shotgun proteomics, ribosome footprinting

## Abstract

Current therapies for neuroblastoma are often ineffective and survivors suffer from severe long-term therapy related side-effects, underscoring the need for identification of novel drugging strategies. We performed an in-depth evaluation of phenotypic and molecular responses following exposure of neuroblastoma cells to the rocaglate CR-1-31-B, scrutinizing its mode-of-action through integrative ribosome footprinting and shotgun proteome profiling. We could show that CR-1-31-B significantly reduces tumor growth *in vivo* without apparent toxicity. By means of combined ribosome footprinting and transcriptome analysis we uncovered that CR-1-31-B treatment downregulates translation efficiencies of several major neuroblastoma dependencies including *MYCN*, *CCND1* and *ALK* as well as factors involved in the G2/M checkpoint. Upregulated targets are enriched for oxidative phosphorylation pathway components and DNA repair. At the proteome level, CR-1-31-B imposed downregulation of a FOXM1 driven signature, including the FOXM1 target gene *TPX2*. We show that neuroblastoma cells are dependent on TPX2 for growth and DNA repair and further demonstrate enhanced CHK1 sensitivity upon TPX2 knockdown. Next, we also observed synergistic effects of CHK1 inhibition with CR-1-31-B. In conclusion, our data support CR-1-31-B as a potent novel therapeutic agent in neuroblastoma, in particular in combination with DNA damage or replication stress inducing agents.

## Introduction

Deregulation of translation is increasingly being recognized as a significant adaptive event during cancer development and progression^1^. Consequently, specific cancer dependencies related to translation have therefore emerged as novel entry points for precision medicine^2^. Translational control is a complex orchestrated process involving ribosomal complexes as well as translation initiation and elongation machineries. The eIF4F translation initiation complex is a heterotrimeric complex composed of a 5’ mRNA cap-binding subunit eIF4E, the large scaffolding protein eIF4G that mediates recruitment of the 40S ribosome subunit and the ATP-dependent RNA helicase eIF4A. Previously, Wolfe *et al*. reported dependency of T-cell leukemia lymphoblasts on inhibition of eIF4A, through the effect on oncogenes such as *MYC* and *NOTCH1*, marked by RNA G-quadruplex structures in their 5’UTR region^3^. Plant-derived rocaglates such as silvestrol, are strong translation inhibitors by targeting the RNA helicase eIF4A. More recently, the therapeutic efficacy of the synthetic rocaglate CR-1-31-B was shown for pancreatic ductal adenocarcinoma, with eIF4A driving the translation of key KRAS signaling pathway components^4^ and concomitant upregulation of glutamine reductive carboxylation installing metabolic reprogramming^5^. Interestingly, a recent report revealed that eIF4A1 is overexpressed in neuroblastoma, a pediatric cancer of the sympathetic nervous system, with CR-1-31-B reducing neuroblastoma cell viability^6^. In this study, we evaluated CR-1-31-B efficacy *in vitro* and *in vivo* in neuroblastoma PDX murine models and explored the underlying mechanism-of-action of CR-1-31-B through a detailed landscaping of the affected translatome by means of integrated ribosome footprinting, shotgun proteomics and RNA sequencing. CR-1-31-B affects several major neuroblastoma dependencies including *MYCN*, *CCND1* and *ALK*, as well as the FOXM1 target *TPX2*. We further demonstrate that TPX2 knockdown or CR-1-31-B drugging sensitizes neuroblastoma cells to pharmacological CHK1 inhibition.

## Results

### CR-1-31-B reduces neuroblastoma cell growth *in vitro*

The eIF4A1 protein is an essential factor for translation initiation, which belongs to an extensive family of DEAD box RNA helicases, with eIF4A1 recently reported to be significantly overexpressed in neuroblastoma tumors compared to non-neoplastic tissue^6,7^. Targeting specific translation factors, that are altered in expression or activity in human cancers, has been shown as promising therapeutic strategy. Therefore, we aimed to evaluate pharmacological eIF4A1 inhibition using the rocaglate CR-1-31-B in the childhood cancer neuroblastoma^4,5^. Recent data shows that CR-1-31-B negatively affects *in vitro* cell growth of SH-SY-5Y and Kelly neuroblastoma cells^6^. We evaluated the phenotypic response to CR-1-31-B exposure in an extended neuroblastoma cell line panel in a dose range over 72h. By means of CellTiter-Glo analysis, we could show that cell viability was negatively affected following 72h of CR-1-31-B exposure, with similar low nanomolar IC50 values achieved across the tested cell line panel (**Figure 1A**). In addition, the negative impact of CR-1-31-B on neuroblastoma cell viability was also demonstrated in two different neuroblastoma spheroid models; NB039 (*MYCN* amplified) and AMC772T (*MYCN* non-amplified) following 120h of drug exposure (**Figure 1B**), achieving similar low nanomolar IC50 values as obtained in the cell line panel.

**Figure 1:**
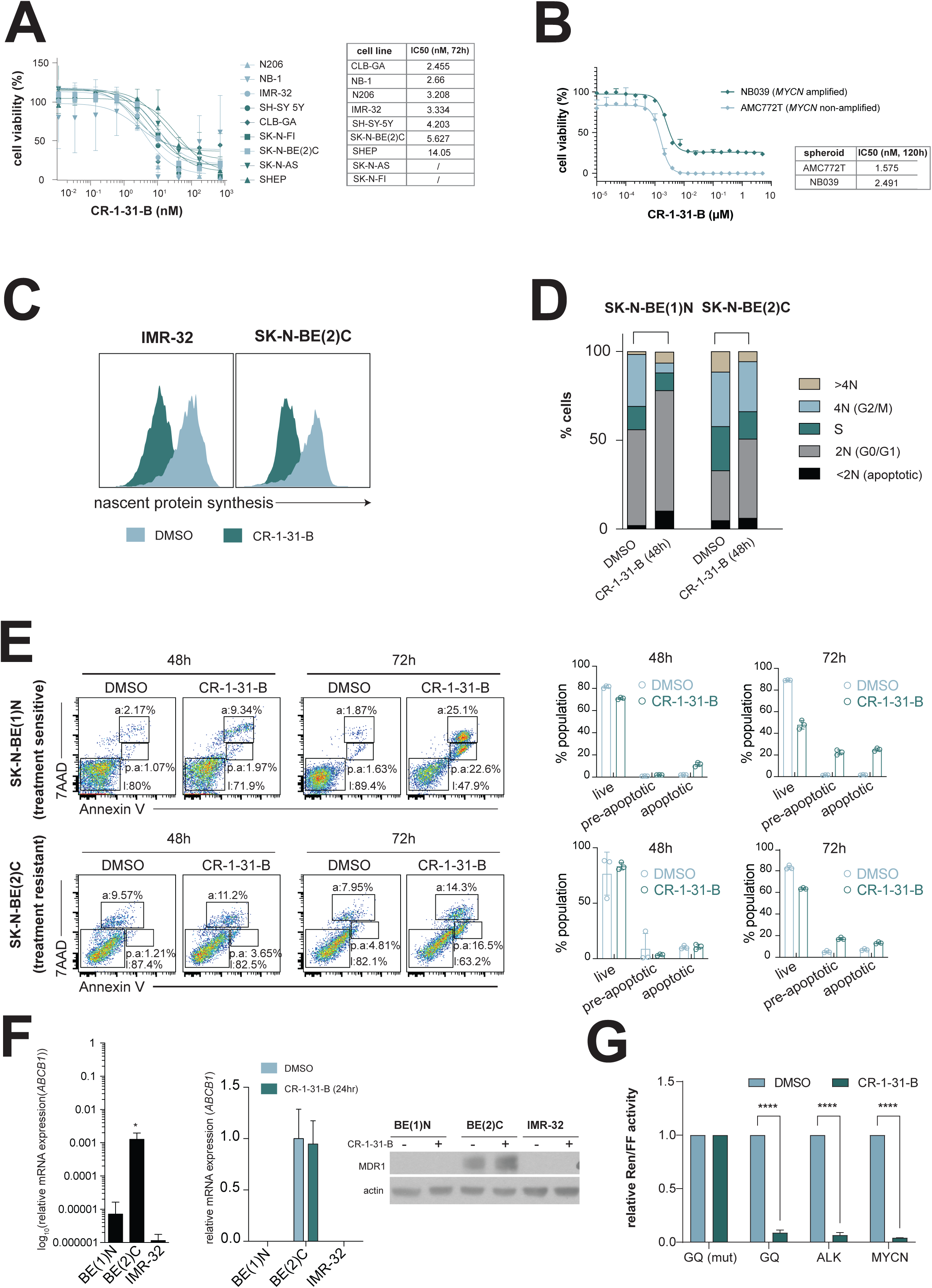
The rocaglate CR-1-31-B reduces neuroblastoma cell growth *in vitro* and affects eIF4A driven translation of *MYCN* and *ALK*. **(A)** Representative plot displaying cell viability measurements across a neuroblastoma cell line panel following exposure (72h) to CR-1-31-B; (n=3) **(B)** Cell viability measurements in two different neuroblastoma spheroid models (120h); **(C)** Histograms showing the change in fluorescent intensity (indicating nascent protein synthesis) of cell lines [IMR-32 (*left*) and SK-N-BE(2)C (*right*)] treated with DMSO (*blue*) or CR-1-31-B (25 nM) (*green*) for 24h; **(D)** Stacked bar graphs showing cell cycle phases in SK-N-BE(1)N and SK-N-BE(2)C neuroblastoma cell lines treated with either DMSO or CR-1-31-B for 48h; **(E)** Flow cytometry analysis of 7AAD and AnnexinV in SK-N-BE(1)N and SK-N-BE(2)C neuroblastoma cell lines treated with DMSO or CR-1-31-B (10nM) for 48h and 72h with (*left*) representative scatter plots representing the live, pre-apoptotic or apoptotic cell populations and (*right)* bar graphs showing the cytotoxic effects of CR-1-31-B (n=3); **(F)** Expression of the MDR1 drug efflux pump at the mRNA level (*ABCB1* gene) in SK-N-BE(1)N, SK-N-BE(2)C and IMR-32 cells under default conditions (*left*) or upon CR-1-31-B treatment (24h, 25 nM) (*middle*) and at protein level (*right*) **(G)** Bar graphs demonstrating the relative luciferase activity (Renilla (Ren)/Firefly luciferase (FF)) of the 5’UTR of *ALK* and *MYCN* in response to DMSO or CR-1-31-B treatment. Technical replicates n=3, biological replicates n=4, 2-tailed student’s t-test, P<0.001***.

To confirm the inhibition of protein translation by CR-1-31-B, nascent protein synthesis was measured 24h post-treatment with 25 nM of CR-1-31-B using metabolic labelling with L-azidohomoalanine (AHA). Indeed, novel protein synthesis was decreased in both IMR-32 and SK-N-BE(2)C neuroblastoma cells in comparison to control treatment (DMSO) (**Figure 1C**). In a next step, we aimed to analyze the impact of CR-1-31-B treatment on cell cycle distribution and compared the response of a neuroblastoma cell line derived from a patient at diagnosis (SK-N-BE(1)N) versus the matched cell line derived from that same patient post-treatment (SK-N-BE(2)C). Interestingly, cell cycle analysis demonstrated a reduction in G2/M phase in SK-N-BE(1) cells, whereas this could no longer be observed in the post-treatment SK-N-BE(2)C cells (**Figure 1D**). Further, induction of cell death was measured by flow cytometry using Annexin V-7AAD staining in the same cell line pair after 48h and 72h treatment with either DMSO or CR-1-31-B. At 48h post-treatment, a slight increase in cell death (pre-apoptosis and apoptosis) was observed in SK-N-BE(1)N while SK-N-BE(2)C cells showed no increase in cell death. Further, cell death increased up to 50% in SK-N-BE(1)N cell line after 72 hours with CR-1-31-B treatment, while only increasing up to 25% in SK-N-BE(2)C cells (**Figure 1E**). To explain these differences in *in vitro* drug response to CR-1-31-B, we hypothesized that this could be linked to P-glycoprotein (encoded by *ABCB1*) drug efflux pump expression, given that previous reports indicated rocaglates like CR-1-31-B as a substrate^8,9^. Indeed, treatment sensitive SK-N-BE(1)N and IMR-32 cells displayed much lower *ABCB1* mRNA expression (**Figure 1F**, *left*) and according MDR1 protein expression (**Figure 1F**, *right*) compared to post-treatment SK-N-BE(2)C neuroblastoma cells, with MDR1 expression not affected by CR-1-31-B treatment (**Figure 1F**, *middle*). Taken together, we show that CR-1-31-B is a potent drug targeting most of the tested neuroblastoma cell lines but also reveal that neuroblastoma cells exhibiting enhanced drug efflux pump expression exhibit reduced drug sensitivity.

### CR-1-31-B targets the translation of the two major neuroblastoma oncogenes, *MYCN* and *ALK*

Given that the translation of genes with 5’UTR containing G-quadruplex (further denoted as GQ) sequences reduces in response to CR-1-31-B treatment^3^, we applied a dual-luciferase reporter assay for 5’UTR regions of the two major neuroblastoma oncogenes, *MYCN* and *ALK*, both containing GQs in their promotor regions. Data was normalized by read outs from HEK293 cells infected with the firefly luciferase plasmid (pGL4.13). We observed significantly reduced reporter signal following CR-1-31-B exposure in comparison to DMSO treatment for both assays supporting an impact of drugging on translation of both genes. To ensure this inhibition is due to presence of a functional GQ sequence, we also included cells transfected with a GQ sequence containing reporter (positive control) and a mutant GQ reporter (negative control). As expected, the functional GQ reporter showed similar downregulation in reporter activity as the *ALK* and *MYCN* reporters following CR-1-31-B treatment. As anticipated, no change in reporter signal could be observed with the mutant GQ reporter following CR-1-31-B treatment (**Figure 1G**). In summary, our data reveal direct effects of CR-1-31-B on translation of *MYCN* and *ALK*, two major neuroblastoma oncogenes, which likely explains at least part of the drug cytotoxicity for the tumor cells.

### CR-1-31-B reduces neuroblastoma cell growth *in vitro*

In a next step, we evaluated the therapeutic efficacy of CR-1-31-B *in vivo* using two different patient-derived xenograft (PDX) murine models, one with *MYCN* amplification (**Figure 2A and 2B, *left***) and one without *MYCN* amplification (**Figure 2A and 2B, *right***). Our results show that only the PDX with *MYCN* amplification is significantly responsive to treatment with CR-1-31-B when compared to the vehicle (5.2%Tween 80 and 2% DMSO) treated animals. In line, H&E staining of the PDX tumors showed the typical appearance of a neuroblastoma tumor, characterized by uniformly sized cells, containing round to oval hyperchromatic nuclei. The number of tumor cells was clearly reduced after CR-1-31-B treatment (**Supplementary Figure 1**). In line with differential sensitivity for CR-1-31-B between SK-N-BE(1)N and SK-N-BE(2)C cells, we hypothesized that the differential sensitivity between the two PDX models could also be linked to a difference in drug efflux pump expression. Indeed, tumor cells derived from the *MYCN* non-amplified PDX model displayed much higher *ABCB1* expression compared to tumor cells of the *MYCN* amplified PDX model (**Figure 2C**).

**Figure 2:**
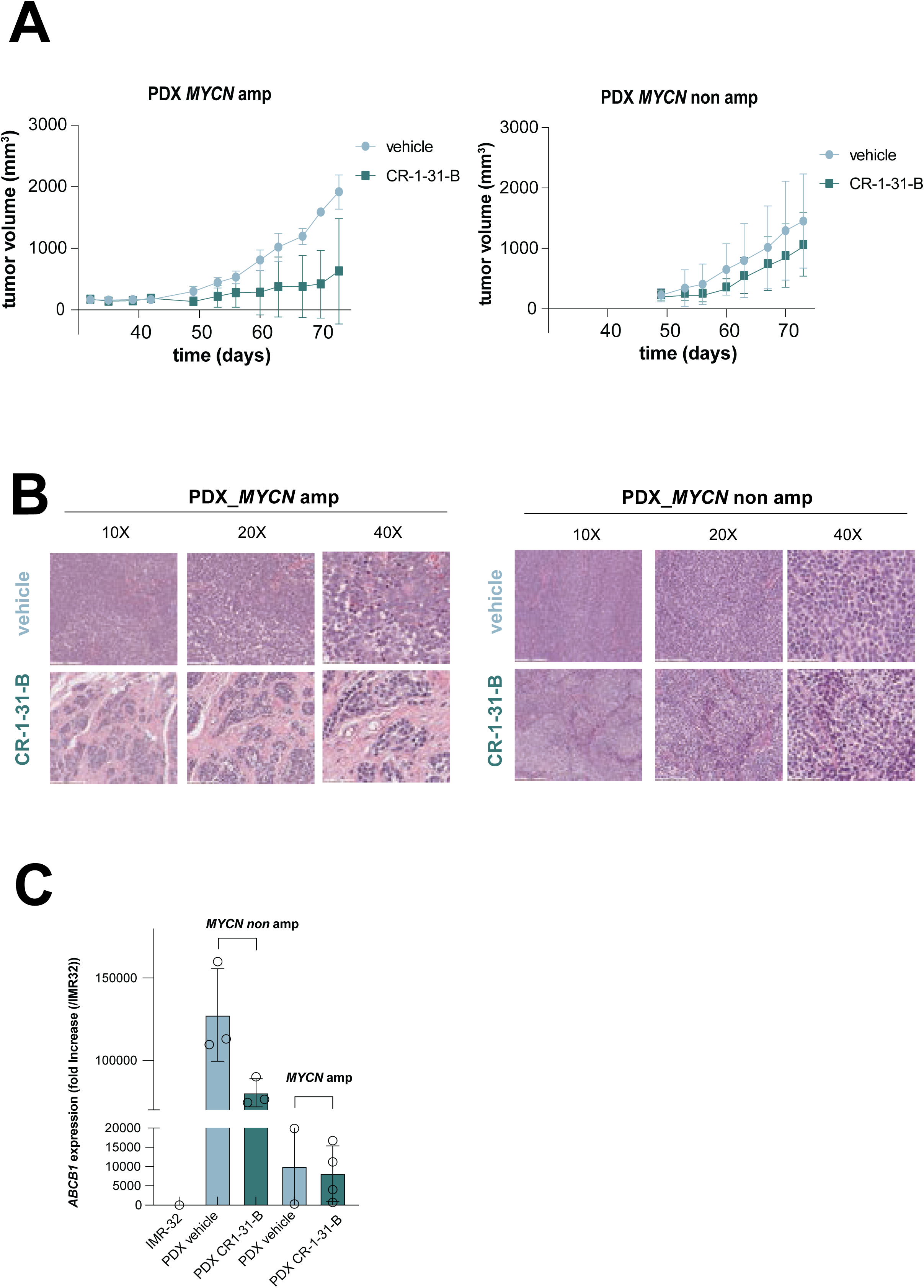
Significant reduction of tumor growth following CR-1-31-B treatment *in vivo* in a patient-derived murine neuroblastoma tumor models. **(A)** Tumor volume (mm^3^) measured over time in two neuroblastoma patient derived xenografts (*MYCN* amplified (*left)* versus non-amplified (*right)*), treated with either vehicle (5.2%Tween 80 and 2% DMSO) or CR-1-31-B (0.25 mg/kg) for 4 weeks; **(B)** Representative H&E sections of neuroblastoma tumor bearing mice treated either with vehicle (5.2%Tween 80 and 2% DMSO) or CR-1-31-B; **(C)** RT-qPCR for *ABCB1* in IMR-32 neuroblastoma cells versus PDX derived tumor cells treated with vehicle or CR-1-31-B. Unpaired t-test with Welch’s correction (two-tailed), P>0.05 ns and P<0.05*.

### Time resolved dissection of the proteome landscape following CR-1-31-B exposure

Following the observation of strong cytotoxic responses to CR-1-31-B exposure both *in vitro* and *in vivo*, we aimed to evaluate the molecular impact of CR-1-31-B treatment on the proteome landscape. To this end, we performed a time course analysis (3h, 6h, 12h, 24h, 48h and 72h) of global protein expression profiles of *MYCN* amplified IMR-32 neuroblastoma cells treated with CR-1-31-B using mass spectrometry-based shotgun proteomics. First, principal component analysis (PCA) analysis showed that the treated samples up to 12h of CR-1-31-B exposure cluster together with the DMSO control samples (**Figure 3A**), implicating that these time points are too early to identify differential protein expression compared to controls. In contrast, the samples from 24h, 48h and 72h under treatment clearly clustered separate from the control treated samples (**Figure 3B**) and revealed in total 271 proteins with significant expression difference over time (from 24h to 72h post-treatment).

**Figure 3:**
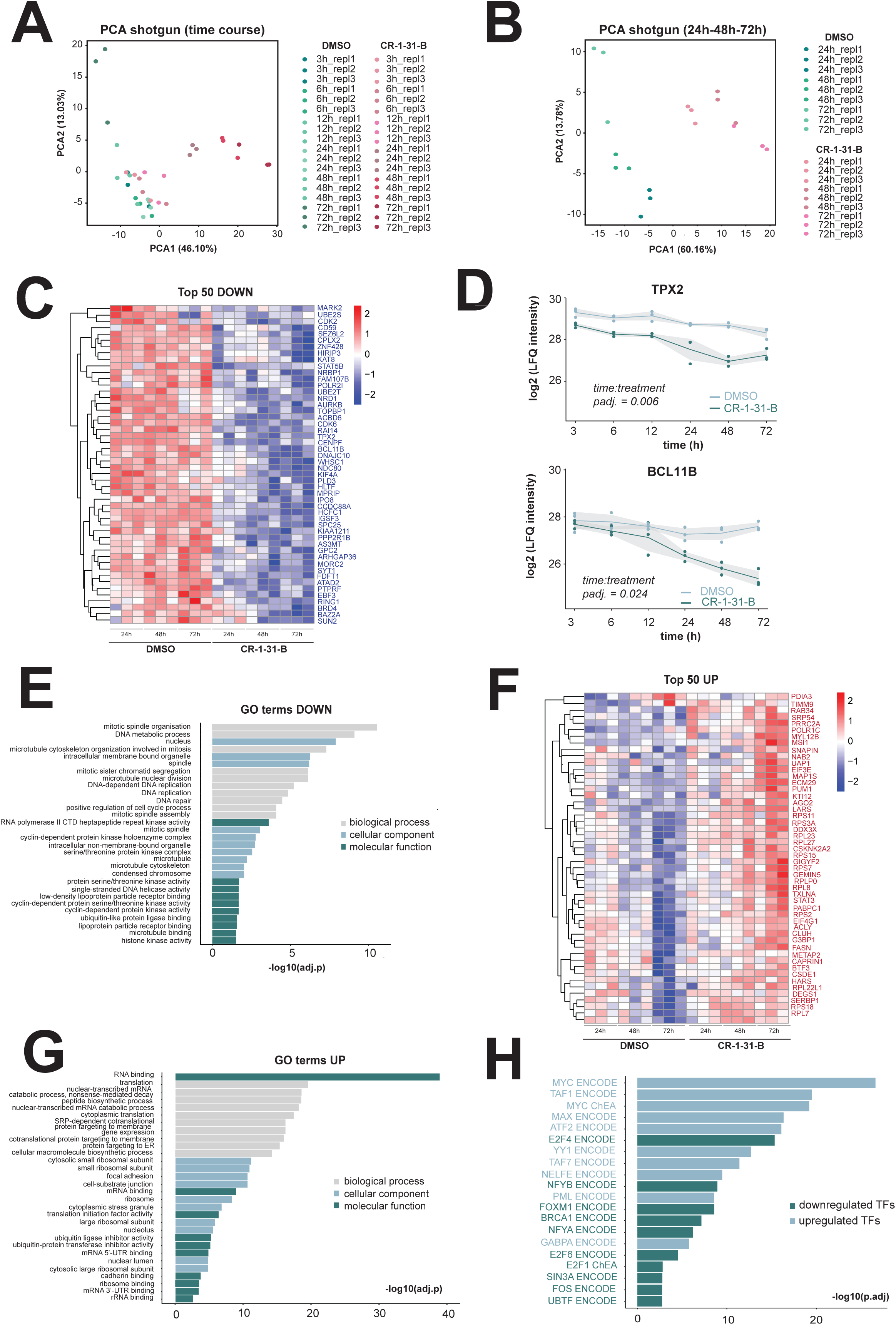
Shotgun proteome profiling following long-term CR-1-31-B treatment of IMR-32 cells. **(A)** Principal Component Analysis (PCA) for all samples included in the shotgun proteome profiling of IMR-32 cells treated with either DMSO or CR-1-31-B; **(B)** PCA analysis for a subset of samples of the shotgun proteome profiling (24h-48h-72h); **(C)** Heatmaps displaying the top 50 downregulated proteins upon CR-1-31-B treatment of IMR-32 cells in a timeseries experiment (24h, 48h, 72h); **(D)** Log_2_(LFQ) plots for the proteins TPX2 and BCL11B in this timeseries; **(E)** Gene ontology analysis for proteins significantly downregulated upon CR-1-31-B treatment of IMR-32 cells in this timeseries; **(F)** Heatmaps displaying the top 50 upregulated proteins upon CR-1-31-B treatment of IMR-32 cells in a timeseries experiment (24h, 48h, 72h); **(G)** Gene ontology analysis for proteins significantly upregulated upon CR-1-31-B treatment of IMR-32 cells in this timeseries experiment; **(H)** ChEA enrichment (EnrichR: maayanlab.cloud/Enrichr/) analysis for the set of downregulated proteins upon CR-1-31-B treatment in IMR-32 cells in this timeseries.

The list of downregulated proteins by CR-1-31-B treatment includes several genes involved in DNA replication (TPX2, NDC80, AURKB, CDK2/6, ATAD2 and CENPF) and DNA damage repair (BCL11B, TOPBP1, HLTF and SUN2)(**Figure 3C and 3D),** as further confirmed by gene ontology analysis for the list of downregulated proteins (**Figure 3E**). The set of upregulated proteins following CR-1-31-B treatment is enriched for ribosomal proteins and factors involved in translation regulation (**Figure 3F and 3G**), potentially as compensatory effect for CR-1-31-B as translation inhibitor. Enrichment analysis for transcription factors (ChEA module, EnrichR) that could regulate the expression of genes part of these signatures, revealed enrichment for E2F4-FOXM1 and BRCA1 targets for the set of CR-1-31-B downregulated factors (**Figure 3H**). We previously identified FOXM1 as key transcriptional regulator in neuroblastoma^10,11^, regulating the expression of important genes required for G2/M transition and DNA repair/replication^12^. Notably, we identified MYC amongst the putative transcriptional regulators of the factors upregulated following CR-1-31-B treatment. Given that this set was primarily enriched for translation initiation factors as well as ribosomal protein components is suggestive for the induction of a translation compensatory mechanism imposed by MYC, in line with previous reports clearly establishing the role of MYC in driving protein translation.

### Short-term exposure to CR-1-31-B affects translation efficiencies of neuroblastoma dependency genes including *CCND1* and *SOX11*

We anticipated that CR-1-31-B likely affects translational efficiencies for regulated targets prior to being detectable at protein expression levels. Given that protein expression changes were only prominently observed from the 24h timepoint on, we therefore evaluated earlier molecular effects of CR-1-31-B treatment in terms of translation efficiency differences through combined ribosome profiling and RNA-sequencing after 6h of drug exposure. The deltaTE method^13^ was used to integrate changes in mRNA and ribosome protected fragments (RPF) levels and allowed for further subcategorization of transcripts into exclusive, intensified, buffered (special) or forwarded categories (**Figure 4A and Supplementary Table S1**). The ‘exclusive’ category (**Figure 4A**, *top left*), indicates significant changes in RPF that are not driven by transcriptional changes. The ‘intensified’ class (**Figure 4A**, *top right*) refers to the changes in translation efficiencies that are driven by both RPF and transcription. The ‘buffered (special) category’ (**Figure 4A**, *bottom left*) reports changes in translation efficiency enhanced by transcriptional changes that are not RPF driven. Finally, the class of forwarded genes (**Figure 4A**, *bottom right*) are driven by changes in transcription. Notably, Gene Ontology (GO) analysis for the downregulated ‘exclusive’ hits revealed a significant enrichment for mitotic spindle and PI3K/AKT signaling (**Figure 4B**, *top left*) and upregulated target enrichment for oxidative phosphorylation and DNA repair (**Figure 4B**, *top right*). Next, GO analysis for the ‘buffered (special)’ category revealed significant downregulation of G2/M checkpoint targets (**Figure 4B**, *bottom left*) and upregulation of DNA repair (**Figure 4B**, *bottom right*). In each of these classes, we scrutinized the transcripts differentially regulated by CR-1-31-B for the presence of a GQ pattern in their 5’UTR region as previously described^14^. For this analysis, the background set was defined by the transcripts for which the translation efficiencies were not significantly altered upon CR-1-31-B treatment. We identified that 2020 out of the 8971 background genes (18.40%) had at least 1 GQ in their 5’UTR region, while each of the defined transcript classes displayed higher enrichment for GQ enriched sequences: 22% in the buffered special, 24.90% in the exclusive class, 28% for the forwarded class and 35.10% in the intensified class of transcripts. In addition, transcripts that contained GQ sequences in each of the classes were enriched in the set of downregulated targets following CR-1-31-B compared to the upregulated targets (**Figure 4C**). Moreover, we evaluated the mean 5’UTR length of each GQ containing target differentially regulated by CR-1-31-B and we could show that each of the defined classes was enriched for genes with longer 5’UTR regions compared to background (**Figure 4D**). The mean 5’UTR length for the background was 310.15 bp while for the ‘buffered special’ it was 355.79 bp, 311.44 bp for the ‘exclusive’ class, 338.86 bp for the ‘forwarded’ class and 350.26 bp for the ‘intensified’ class. Next, we performed complementary transcription factor motif enrichment analysis using ‘Simple Enrichment Analysis of motifs’ (SEA) and identified the ‘Myc-associated zinc finger protein’ (MAZ) motif to be represented in 44% of all sequences (p-value: 1.47e-28) (**Figure 4E**), indicating that MYC(N) is a key putative regulator of GQ enriched targets. Within the ‘exclusive’ gene class, we identified *CCND1*^15^ (**Figure 4F**, *left*), a key neuroblastoma oncogene, and within the ‘buffered special’ category amongst others *SOX11*^16^ (**Figure 4F**, *middle*) with significantly downregulated translation efficiencies upon CR-1-31-B treatment. Interestingly, in line with our GQ reporter assay (**Figure 1F**), we could confirm that the translation efficiency of *ALK* was indeed strongly downregulated upon CR-1-31-B treatment **(Figure 4F**, *right*).

**Figure 4:**
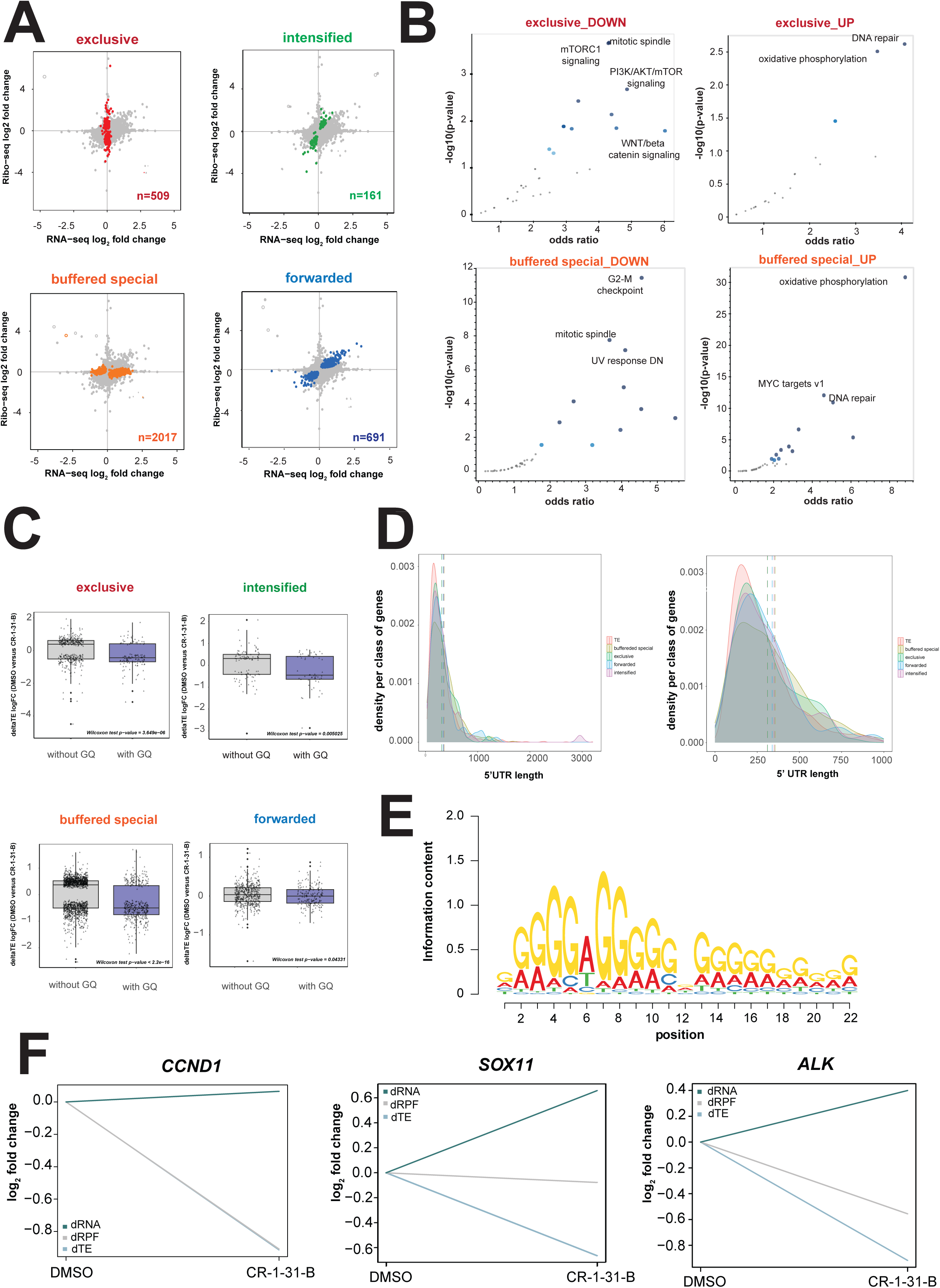
Landscaping translatome changes following short-term CR-1-31-B exposure through integrated ribosome profiling and RNA-sequencing. **(A)** CR-1-31-B imposed translatome changes were further subclassified using the deltaTE method^13^; **(B)** Gene Ontology (GO) for up- and downregulated transcripts within the ‘exclusive’ and ‘buffered special’ category; **(C)** Boxplots showing the log fold changes in translational efficiencies for DMSO versus CR-1-31-B treatment for targets with our without GQ motifs in their 5’UTR region in the categories as defined in *panel A*; **(D)** Distribution plots showing the mean 5’UTR length of transcripts in each of the categories as defined in *panel A* compared to background; **(E)** Simple Enrichment Analysis (SEA) revealed the MAZ motif as most prominent sequence motif within the 5’ UTR region of the different transcripts enriched with a GQ motif; **(F)** Translation efficiencies of neuroblastoma oncogenes *CCND1*, *SOX11* and *ALK* are downregulated upon CR-1-31-B exposure;

### TPX2 is a CR-1-31-B sensitive dependency gene in neuroblastoma

Our integrated ribosome footprinting, transcriptome, and shotgun proteome profiling analyses revealed the broad molecular impact of CR-1-31-B treatment. In addition to the impact on key targets such as *MYCN*, *ALK* and *CCND1*, we next decided to identify possible additional targets contributing to the observed drug response. To this end, we selected all proteins across the entire time series (3h to 72h) with significant differential expression for the interaction between both the time and treatment component. From this hit list, we subsequently calculated the mean difference in LFQ intensities between DMSO versus CR-1-31-B treatment across this time series as a ranking parameter for dynamic target regulation. This resulted in a final ranking from which we selected the top 50 significantly downregulated proteins by CR-1-31-B to identify a candidate target downstream of CR-1-31-B for further functional evaluation (**Figure 5A**). Subsequently, expression of these 50 targets was evaluated in a cohort of primary versus relapsed neuroblastoma cases^17^ (GSE16476) and we could show that 18 out of 50 genes were significantly higher expressed in relapsed versus primary neuroblastoma, with TPX2 as a one of the top ranked genes (**Figure 5B**). This further underscores the putative importance of TPX2 in neuroblastoma tumor biology, in line with previous studies showing that high *TPX2* expression significantly correlates with poor neuroblastoma patient survival^18^. Moreover, we previously dissected the time-resolved transcriptome profiles of developing tumors in a TH-MYCN driven murine neuroblastoma model^19^. From this dataset, the relevance of TPX2 in murine neuroblastoma tumor formation is further underscored as *Tpx2* expression levels are significantly elevated both in hyperplastic lesions (week 2) and full-blown tumors (week 6) versus ganglia from wildtype mice (**Figure 5C**). As expected for a highly transcribed gene, CUT&RUN mapping of genome-wide binding sites of MYCN in IMR-32 neuroblastoma cells identified TPX2 as a direct MYCN bound target (**Figure 5D**). Furthermore, through immunoblotting we could confirm that TPX2 protein levels are strongly downregulated upon CR-1-31-B exposure versus control treatment (**Figure 5E**). Interestingly, it was previously reported in the context of acute T-cell leukemia that the translational efficiency of TPX2 was significantly downregulated with CR-1-31-B, related to the presence of a 12-mer GC rich motif in its 5’UTR region with the potential of G-quadruplex formation^3^. In that respect, we also performed a GQ reporter assay for *TPX2* as done for both *MYCN* and *ALK* (**Figure 1F**) and confirmed that the reporter activity in the presence of the *TPX2* 5’UTR region is indeed significantly downregulated compared to DMSO treatment to a similar extend as the GQ positive control (**Figure 5F**). In view of the above, we aimed to study in more depth the phenotypic and molecular consequences of deregulated TPX2 expression in neuroblastoma cells. Transient knockdown of TPX2 in IMR-32 cells, as confirmed through RT-qPCR and immunoblotting (**Figure 5G and 5H)** using three independent siRNAs, significantly reduced cell confluence as measured over time by Incucyte live cell imaging (**Figure 5I**). Concomitant induction of apoptosis was shown by significantly increased caspase 3/7 luminescent signal upon TPX2 knockdown compared to controls (**Figure 5J**). Given that TPX2 translation is blocked by CR-1-31-B, we also evaluated the phenotypic impact of combined TPX2 protein downregulation (using siRNAs) with CR-1-31-B drug exposure (TPX2 translation block). Interestingly, growth of IMR-32 neuroblastoma cells could be completely suppressed when combining TPX2 downregulation with CR-1-31-B treatment compared to cells with reduced TPX2 expression treated with DMSO (**Figure 6A**). We could confirm that the confluence of parental IMR-32 neuroblastoma cells was only negatively affected to limited extent as expected upon exposure to low dose CR-1-31-B concentrations (**Supplementary Figure 2A**). Altogether, TPX2 could, in addition to MYCN, CCND1 and ALK, putatively impact on neuroblastoma cell viability following exposure to CR-1-31-B. Further experiments are warranted to explore the exact contribution of each of these (and possibly also other) factors to the observed drug response.

**Figure 5:**
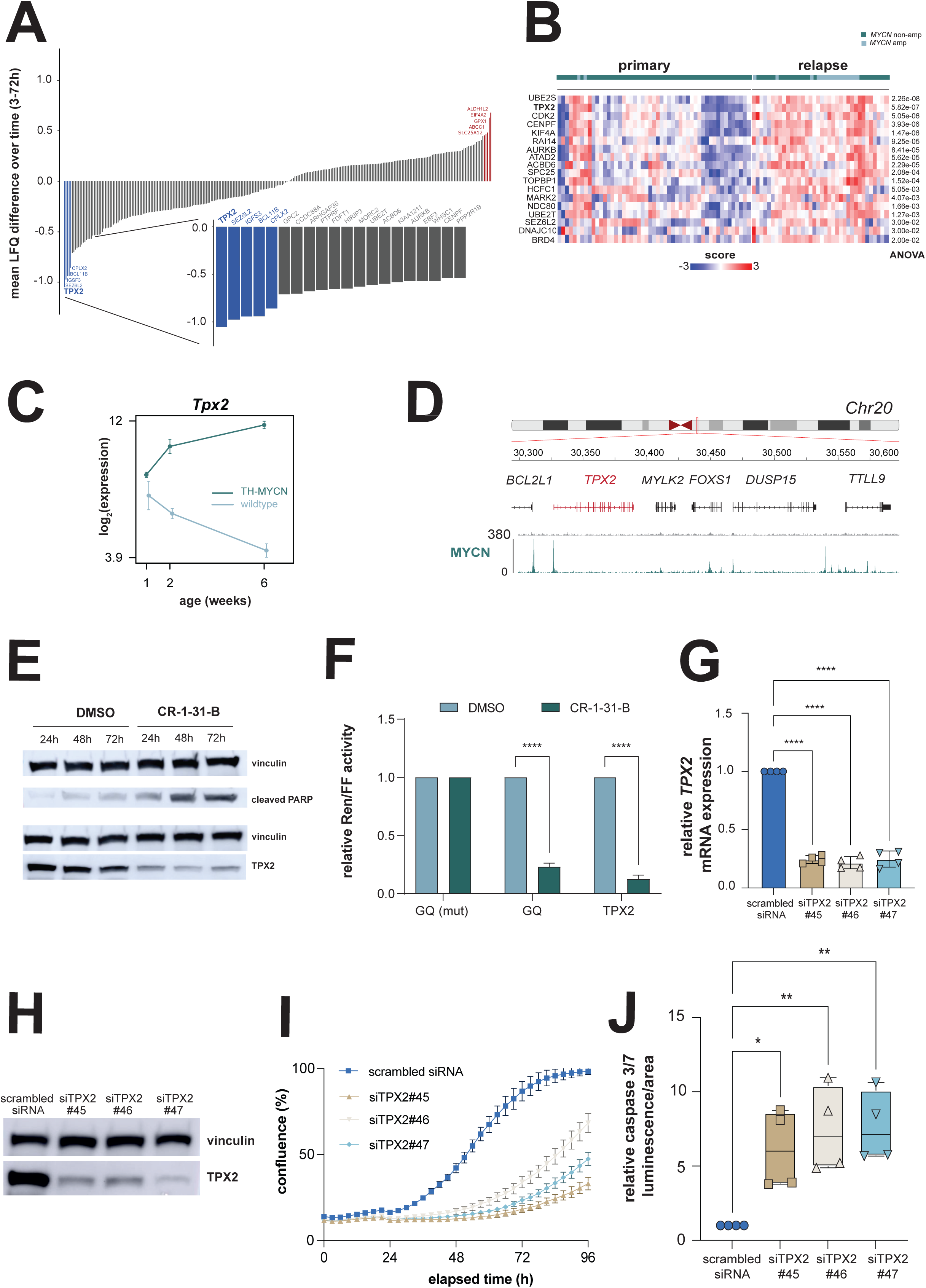
Prioritization of TPX2 as a novel dependency factor modulated by CR-1-31-B in neuroblastoma. **(A)** Ranking of mean LFQ differences of significantly differentially expressed proteins following CR-1-31-B treatment versus DMSO in the shotgun timeseries experiment with a significant p-value for the interaction between the time and treatment components; **(B)** Heatmap displaying the expression of the 18 significantly differentially expressed genes in primary versus relapsed neuroblastomas (GSE16476) out of the top-50 significantly downregulated proteins following CR-1-31-B exposure (*see* Figure 3C); **(C)** Tpx2 expression is significantly upregulated during murine neuroblastoma tumor development both in hyperplastic lesions (*week 2*) and in full-blown tumors (*week 6*) compared to wildtype mice; **(D)** Mapping genome-wide MYCN binding sites by CUT&RUN shows direct binding of MYCN to the *TPX2* promotor region; **(E)** Representative image of immunoblotting for cleaved PARP and TPX2 following CR-1-31-B drug exposure (IC50) of IMR-32 cells (biological replicates n=3); **(F)** Bar graphs demonstrating the relative luciferase activity (Renilla (Ren)/Firefly luciferase (FF)) of the 5’UTR of *TPX2* in response to DMSO or CR-1-31-B treatment. Technical replicates n=6, biological replicates n=3, 2-tailed student’s t-test, P > 0.01 = ns, P<0.001***; **(G)** Knockdown of TPX2 using 3 independent siRNAs in IMR-32 cells as confirmed by RT-qPCR (biological replicates n=4), **(H)** Knockdown of TPX2 using 3 independent siRNAs in IMR-32 cells as confirmed by immunoblotting (representative image of 4 biological replicates) ; **(I)** Representative example of confluence (%) as measured over time by Incucyte live cell imaging of IMR-32 cells transfected either with scrambled or TPX2 targeting siRNAs (error bars resulting from technical replicates n=3, biological replicates n=4); **(J)** Relative caspase 3/7 luminescence/area measurements using the Caspase-Glo assay following transfection of IMR-32 cell with scrambled or TPX2 targeting siRNAs (biological replicates n = 4). Statistical test used was one-way ANOVA with Dunnet’s multiple comparison test, P≤0.05*, P≤0.01**.

**Figure 6:**
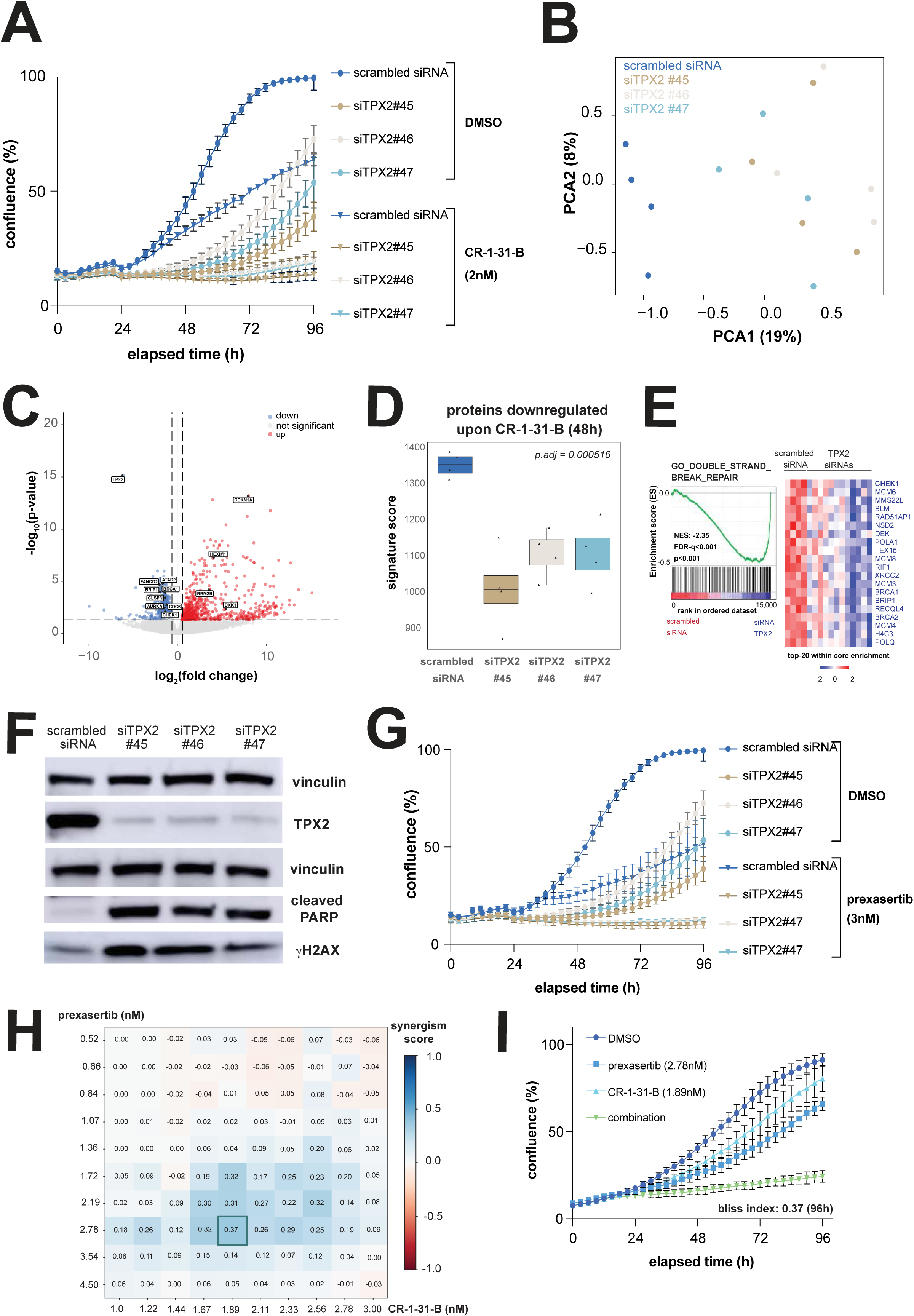
Identification of drug synergism between prexasertib and CR-1-31-B in neuroblastoma. **(A)** Representative example of confluence (%) as measured over time by Incucyte live cell imaging of IMR-32 cells transfected either with scrambled or TPX2 targeting siRNAs combined either with DMSO or CR-1-31-B treatment (error bars resulting from technical replicates n=3, biological replicates n=3); **(B)** Principal Component Analysis (PCA) for all samples included in the transcriptome profiling of IMR-32 cells following transfection with either scrambled or TPX2 targeting siRNAs; **(C)** Volcano plot showing the set of significantly down (*blue*) or upregulated (*red*) genes following transient knockdown of TPX2 in IMR-32 cells; **(D)** Signature score analysis for the set of significantly downregulated proteins in IMR-32 cells 48h post-treatment with CR-1-31-B in the transcriptome profiles of IMR-32 cells with TPX2 knockdown versus controls; **(E)** Gene Set Enrichment analysis shows a strong downregulation of genes involved in DNA double strand break repair following TPX2 knockdown versus controls; **(F)** representative immunoblotting for cleaved PARP and γH2AX upon TPX2 knockdown in IMR-32 cells (biological replicates n=3) ; **(G)** Representative example of confluence (%) as measured over time by Incucyte live cell imaging of IMR-32 cells transfected either with scrambled or TPX2 targeting siRNAs combined either with DMSO or prexasertib treatment (error bars resulting from technical replicates n=3, biological replicates n=3); **(H)** Representative heatmap displaying the Bliss index scores as measured for combined treatment with prexasertib (CHK1 inhibitor) and CR-1-31-B over a large concentration series at 96h post-treatment; **(I)** Representative example of confluence (%) as measured over time by Incucyte live cell imaging of IMR-32 cells upon combined prexasertib and CR-1-31-B treatment with the highest Bliss index score (0.37) over the combined concentration ranges tested as displayed in *panel H* (error bars resulting from technical replicates n=3, biological replicates n=3).

### TPX2 knockdown enhances DNA damage and sensitizes neuroblastoma cells to CHK1 inhibition

Next, we aimed to more profoundly understand the key molecular pathways regulated by TPX2 that take part in this drug response. To this end, we performed transcriptome profiling following TPX2 knockdown, with PCA analysis indicating clear transcriptome differences between TPX2 siRNA versus scrambled siRNA transfected neuroblastoma cells (**Figure 6B**). Genes upregulated following TPX2 knockdown were indicative for the induction of a p53 response gene signature, including *HEXIM1*, *RRM2B*, *CDKN1A* and *DKK1*, with the latter being a presumed tumor suppressor in neuroblastoma^20^. Differential gene expression profiling analysis confirmed significant TPX2 knockdown together with downregulation of genes that play an important role in DNA replication and DNA damage repair in line with the set of downregulated proteins by CR-1-31-B, including *ATAD2*, *AURKA*, *BRIP1*, *FANCD2*, *BRCA1*, *CDC6*, *CLSPN* and *CHEK1* (**Figure 6C and Supplementary Table S2**). The significant overlap between genes downregulated upon TPX2 knockdown with proteins downregulated upon CR-1-31-B exposure could be further confirmed through compilation of all significantly downregulated proteins in IMR-32 cells by CR-1-31-B (versus DMSO controls) 48h post-treatment into a signature of which the activity was quantified as a signature score in IMR-32 cells transfected either with a scrambled or TPX2 targeting siRNA (**Figure 6D**). This analysis indicates that the expression signatures obtained following CR-1-31-B treatment or TPX2 knockdown are significantly overlapping and could therefore explain the combined negative effect on cell confluence (**Figure 6A**). The enrichment for DNA repair pathway genes in the set of downregulated factors following TPX2 knockdown was further confirmed through gene ontology analysis (**Figure 6E**). Notably, recent data highlight a key moonlighting function for TPX2 in concert with AURKA in DNA damage repair and replication fork stability besides its known role in cell cycle regulation^21^. In the process of homologous recombination, it has been reported that knockdown of either TPX2 or AURKA resulted in decreased RAD51 foci^22^. Additionally, Byrum et al. showed that TPX2 regulates BRCA1 recruitment to sites of double stranded breaks^22^. Through immunoblotting we could indeed confirm upregulation of γH2AX as a marker for DNA double strand breaks upon TPX2 knockdown (**Figure 6F**). Interestingly, previous reports indicated that translation inhibition can lead to activation of the ATR-CHK1 pathway^23^ and MYCN induced replication stress has been shown to impose sensitization for DNA damage induction^24^. In the context of neuroblastoma, CHK1 has been reported as a key synthetic lethal target in *MYCN* amplified neuroblastoma, given its role in replication fork stability. These data converged to the hypothesis that CR-1-31-B or TPX2 knockdown could sensitize neuroblastoma cells for pharmacological DNA damage induction. More specifically, we hypothesized that TPX2 downregulation or CR-1-31-B drug exposure could sensitize for CHK1 inhibition using prexasertib and therefore presents a novel entry point for combination therapy. Indeed, neuroblastoma cells with TPX2 downregulation using siRNAs were strongly sensitized for prexasertib treatment compared to control cells (**Figure 6G**), with the confluence of parental cells only to a limited extent affected upon exposure to low dose prexasertib concentrations as expected (**Supplementary Figure 2B**). Moreover, we could show a synergistic drug interaction with low dose CR-1-31-B and prexasertib combination treatment (representative example of 3 biological replicates, *Bliss score: 0.37*) (**Figure 6H and 6I**). Our current data cannot provide detailed mechanistic insights into the contribution of CR-1-31-B mediated TPX2 translational inhibition to the negative impact of CR-1-31-B on neuroblastoma cell viability. Nevertheless, in line with the previously reported role for TPX2 in control of replication stress (besides its canonical role in mitosis), we uncover for the first time a synergistic interaction between TPX2 knockdown and CR-1-31-B drugging with CHK1 pharmacological inhibition, thereby opening novel perspectives for targeted therapy.

## Discussion

Translation is a highly complex biological process and hijacking of translational control is emerging as a common theme in malignant transformation. Most components of the translational machinery have been found to be deregulated in cancer cells. Translational perturbation can boost cancer plasticity through rapid rewiring of the cell proteome while concomitantly reshaping phenotypes. Rocaglates are natural products that inhibit protein synthesis in eukaryotes and have been identified to directly target eIF4F. These molecules have gained significant interest due to their promising pre-clinical activity. The natural rocaglate silvestrol was characterized to bind and block the activity of the eIF4A subunit with nanomolar affinity. The synthetic rocaglate CR-1-31-B inhibits more specifically eIF4A1 and was recently suggested to effectively reduce neuroblastoma cell viability *in vitro*^6^. In this study, we could clearly demonstrate anti-tumor effects of CR-1-31-B *in vivo*. Notably, we observed difference in CR-1-31-B drug sensitivity between the *MYCN* amplified versus *MYCN* non-amplified PDX models, while no significant differences in IC50 values were noted for the *in vitro* screened cell lines. There are several possible reasons that could explain the apparent difference between the data obtained *in vitro* versus *in vivo*. First, most neuroblastoma cell lines are *MYCN* amplified and this study highlights that the translation efficiency of MYCN is affected by CR-1-31-B. Second, very few cell lines exist that faithfully recapitulate the biology of the high-risk *MYCN* non-amplified tumors, hampering a broader evaluation *in vitro*. In this case, PDXs can offer a good alternative to overcome this problem and further *in vivo* experiments will be required to define better the exact determinants of differential drug sensitivity.

Interestingly, one possible explanation could be linked to CR-1-31-B being a substrate for P-glycoprotein drug efflux pumps. Indeed, we showed that IMR-32 and SK-N-BE(1)N neuroblastoma cells, which are sensitive to CR-1-31-B, display low *ABCB1* expression (mRNA level) with no expression of the MDR1 protein. This was in contrast to drug efflux pump expression in the treatment resistant SK-N-BE(2)C cells, which display a higher half maximum inhibitory concentration for CR-1-31-B compared to IMR-32 and SK-N-BE(1)N. Further investigations will be required to explore additional possible factors, including p53 or other mutations, that impact on differential drug sensitivity observed between SK-N-BE(1)N versus SK-N-BE(2)C or the *MYCN* amplified versus the *MYCN* non-amplified neuroblastoma PDX models.

In this study, we performed for the first time an in-depth analysis of both proteome and translation efficiency induced changes upon exposure of neuroblastoma cells to CR-1-31-B. Notably, protein expression differences emerged from 24h post-treatment onwards. While the set of downregulated proteins by CR-1-31-B are enriched for a FOXM1-E2F signature, we observed that the set of upregulated proteins was rather MYC(N) driven. This could relate to the previously established role of MYC(N) as a driver of protein translation. In contrast to the proteome dataset, we could already clearly observe significant changes in translation efficiencies early after treatment (6h), including established oncogenic factors in neuroblastoma such as *CCND1, MYCN* and *ALK*.

Within the set of targets of which the translational efficiencies were downregulated with CR-1-31-B, we could show an enrichment for transcripts with longer 5’UTR sequences enriched with GQs compared to background, which is in line with the recent findings of Volegova *et al.*^25^. Next, we identified TPX2 as another key dependency gene affected by CR-1-31-B. We present for the first time the functional implication of TPX2 downstream of eIF4A in neuroblastoma cells. The TPX2 protein is overexpressed in multiple solid tumor types and besides the canonical role of TPX2 in cell division, moonlighting functions in DNA damage response and replication stress together with AURKA are emerging thus warranting further investigation. Interestingly, translation inhibition has been previously reported to activate the ATR-CHK1 axis and impose upregulation of a replicative stress gene signature^23^. This is in line with our findings that TPX2 knockdown or CR-1-31-B can sensitize neuroblastoma cells to pharmacological CHK1 inhibition. Neuroblastoma has been identified as one of the most highly addicted tumor entities to CHK1 activity and several studies, including from our lab, showed strong potential of this tumor addiction for further translational exploration for combinatorial drugging strategies.

Taken together, we present for the first time a detailed and comprehensive dissection of the molecular signatures following pharmacological eIF4A inhibition in neuroblastoma cells and present amongst others TPX2 as key downstream CR-1-31-B target to putatively protect rapidly dividing cells from DNA damage and replicative stress insults. This could serve as entry point to define rational combination therapies with CR-1-31-B, including novel CHK1 inhibitors, other DNA damage inducing agents or selective TPX2-AURKA protein-protein interaction disruptors such as CAM2602^26^.

## Acknowledgements

We would like to thank prof. dr. Frank Speleman for the guidance and support in the experimental design and conceptualization of this manuscript. We would like to thank Javier Otero and Man Jiang for their technical assistance with *the in vivo* experiments. This research was supported by the Villa Joep Foundation (STI.DIV.2021.0002.01), the Research Foundation Flanders (1197617N and FWO.3F0.2020.0048.01), Olivia Fund, GOA (Ghent University, Belgium, BOF.GOA.2022.0003.03) and the Belgian ‘Kom op Tegen Kanker’ foundation (STI.VLK.2022.0003.01).

## Author contributions

Conceptualization, G.H.W., K.D.; Methodology, C.N., S.L.B., S.P., E.S., L.P. E.D.S., E.H., G.H.W., K.D.; Formal analysis, C.N., S.L.B., S.P., I.N., P.Z., A.S., E.S., L.P., E.H., G.H.W. and K.D.; Investigation: C.N., S.L.B., S.P., I.N., F.M., S.T., F.D.V., A.E, E.S., L.P. and S.G.; Writing – Original Draft: C.N., S.L.B., S.P., I.N., G.H.W., L.P. and K.D.; Writing – Reviewing & Editing, G.H.W. and K.D.; Visualization: C.N., S.L.B., S.P., L.P. and I.N.; Supervision; G.H.W. and K.D., Funding acquisition: G.H.W. and K.D.

## STAR Methods

### RESOURCE AVAILABILITY

#### Lead contact

Further information and requests for resources and reagents should be directed to and will be fulfilled by the lead contact, Kaat Durinck.

#### Materials availability

Plasmids generated in this study are available from the lead contact upon request.

#### Data and code availability

- RNA-seq, RIBO-seq and Cut & Run data have been deposited at GEO and are publicly available as of the date of publication. Proteomics data are available via the ProteomeXchange. Accession numbers are listed in the key resources table. Original western blot images have been deposited at Mendeley and are publicly available as of the date of publication. The DOI is listed in the key resources table. Microscopy data reported in this paper will be shared by the lead contact upon request.
- Original western blot images have been deposited at Mendeley and are publicly available as of the date of publication. The DOI is listed in the key resources table.
- All scripts used in these analyses are available on GitHub: https://github.com/PPOLLabGhent/CR-1-31-B_TranslationInsights
- Any additional information required to reanalyze the data reported in this paper is available from the lead contact upon request.

### EXPERIMENTAL MODEL AND SUBJECT DETAILS

#### Cell lines

- SK-N-AS, female
- SH-SY5Y, female
- SK-N-BE(1)-N, male
- SK-N-BE(2)-C, male
- IMR-32, male
- CLB-GA, male
- NB-1, male
- SH-EP, female
- SK-N-FI, male
- N206, female

See also **Supplementary Table S3.**

Cells were grown in RPMI 1640 medium supplemented with 10% fetal calf serum (FCS), 100 IU/mL penicillin/streptomycin and 2mM L-glutamine at 37°C with 5% CO_2_. Short Tandem Repeat (STR) genotyping was used to validate cell line authenticity prior to performing the described experiments and Mycoplasma testing was done on a monthly basis. Genotyping data of the cell lines have been deposited at Mendeley and are publicly available as of the date of publication. The DOI is listed in the key resources table.

#### Spheroids

- NB039
- AMC772T

Patient-derived neuroblastoma tumour spheroids were grown in DMEM-GlutaMAX medium containing low glucose and supplemented with 25% (v/v) Ham’s F-12 Nutrient Mixture, B27 Supplement minus vitamin A, N-2 Supplement (100X), 100 U/mL penicillin, 100 µg/mL streptomycin, 20 ng/mL epidermal growth factor (EGF) and 40 ng/mL fibroblast growth factor-basic (FGF-2), 200 ng/mL insulin-like growth factor-1 (IGF-1), 10 ng/mL platelet-derived growth factor-AA (PDGF-AA) and 10 ng/mL platelet-derived growth factor-BB (PDGF-BB) at 37°C with 5% CO_2_.

#### Mice

Six to eight weeks old female NOD/SCID/IL2Ry^-/-^ gamma (NSG) mice were purchased (Jackson Laboratory, Bar Harbor, ME) to establish patient derived xenograft models (PDX) as described in the METHOD DETAILS section. Female NOD/SCID/IL2Ry^-/-^ (NSG) mice of six to eight weeks were used for the in vivo treatment as described in the METHOD DETAILS section.

### METHOD DETAILS

#### Cell viability measurements for single compound treatment

The adherent cell lines were plated in 96-well plates at density of 2 × 10^3^ – 1.5 × 10^4^ cells per well, depending on the cell line. Cells were allowed to adhere overnight, after which CR-1-31-B was added in a range of concentrations. Cytotoxicity assays were performed at 48 and 72 hours after treatment with CellTiter-Glo® reagent (Promega). The protocol was adapted, adding 50 μL of reagent at each condition. The results were normalized to the 0.1% DMSO control condition and the different inhibitory concentration values were computed through the GraphPad Prism Software (version 10.0.1). The dose-response curve analysis was performed through ECanything equation assuming a standard slope of -1.0. The error bars in figures represent the standard deviation from three biological replicates.

#### Organoids Cell Viability Screening

Patient-derived neuroblastoma tumour organoids were harvested using Accutase® solution (Sigma-Aldrich), made single cell, filtered using a 70 µm nylon cell strainer (Falcon) and resuspended in appropriate growth medium. Subsequently, cells were plated at densities ranging from 1 × 10^3^ – 6 × 10^3^ cells per well using the Multi-drop™ Combi Reagent Dispenser on repellent black 384-well plates (Corning). Following 24 hours of recovery time, cells were treated with 0.01-5000 nM CR-1-31-B or DMSO (negative control) using the Tecan D300e Digital Dispense (HP). Two technical replicates were included in each experiment and two biological replicates were completed for each patient-derived neuroblastoma tumour organoid. After five days of treatment, ATP levels were measured using CellTiter-Glo 3D® (Promega) according to the manufacturer’s instructions. The results were normalized to the 0.1% DMSO control and data were analysed with GraphPad Prism (version 10.0.1).

#### Nascent protein synthesis assay

1 × 10^6^ Cells were treated with CR-1-31-B (25nM) or DMSO for 22 hours. Then, cells were starved using RPMI methionine-free medium for one hour and incubated with Click-iT AHA (L-azidohomoalanine) metabolic labelling reagent for another hour. Cells were fixed with 4% paraformaldehyde for 15 min and permeabilized with 0.25% Triton-X for 15 min as instructed by Click-iT Cell reaction Buffer Kit. Then, the samples were stained with Alexa Fluor 488 Alkyne Changes in mean florescence intensity of AF488 was measured by Guava Easy-Cyte flow cytometer.

#### Cell Death Assay

Cells were stained with APC Annexin V and 7-AAD as recommended by the manufacturer. In brief, cells were treated with CR-1-31-B (25nM) or DMSO for 24h, 48h, and 72h and washed twice with cell staining buffer and resuspended in 100μL of Annexin V binding buffer. The cells were stained with 1μL of APC Annexin V and 1μL of 7AAD for 15 min. 500μL of Annexin V binding buffer was added to each sample. Analysis was performed with a BD LSR Fortessa cell analyzer and FlowJo software 10.4.1 (Tree Star).

#### Cell cycle analysis

Cell cycle analysis was performed using Guava cell cycle reagent (Part number 4500-0220) as instructed by manufacturer. In brief, to induce cell cycle arrest and synchronize the cells, 1 × 10^6^ cells were cultured in serum-free media for 24h. The cells were treated with CR-1-31-B (25nM) or DMSO for 24h, 48h, and 72h and fixed with 70% ethanol for 1 hour. Afterwards, cells were washed and incubated with 200μL of Guava cell cycle reagent at RT for 30 min in the dark. The guava easyCyte flow cytometer was used to analyze the cell cycle phases.

#### Dual luciferase reporter assay

The 5’UTR of ALK [F primer: AGCTaagcttGGGGGCGGCAGCGGT, R primer: AATTaagcttCCCGCCGGAGGAGGC], 5’UTR of N-MYC [F primer: agctAAGCTTgtcatctgtctggac, R primer: aattAAGCTTcggctcgcctcccgg] and the controls for this experiment, GQ or mutant GQ sequence (the sequence published previously^3^) were cloned separately in Renilla luciferase pGL4.73 plasmids as previously shown^3^. Empty firefly luciferase pGL4.13 plasmid was used as internal controls. HEK293 cells were infected with either of above pGL4.73 plasmids and empty firefly luciferase pGL4.13 plasmid. 5 × 10^5^ cells were treated with either DMSO or CR-1-31-B (25nM) for 24 hours. Dual-luciferase reporter assay was used to lyse the cells and read the luciferase activity as instructed by manufacturer. The signal was measured using Luminometer.

The 5’UTR of tpx-2 was ordered as G-block and cloned into pGL4.73 plasmids using Gibson assembly:taaagccaccagtggactcacgcaggcgcaggagactacacttcccaggaactccgggccgcgttgttcgctggtacctccttctgacttccggtattgctgcggtctgtagggccaatcgggagcctggaattgctttcccggcgctctgattggtgcattcgactaggctgcctgggttcaaaatttcaacgatactgaatgagtcccgcggcgggttggctcgcgcttcgttgtcagatctgaggcgaggctaggtgagccgtgggaagaaaagagggagcagctagggcgcgggtctccctcctcccggagtttggaacggctgaagttcaccttccagcccctagcgccgttcgcgccgctaggcctggcttctgaggcggttgcggtgctcggtcgccgcctaggcggggcagggtgcgagcaggggcttcgggccacgcttctcttggcgacaggattttgctgtgaagtccgtccgggaaacggaggaaaaaaagagttgcgggaggctgtcggctaataacggttcttgatacatatttgccagacttcaagatttcagaaaaggggtgaaagagaagattgcaactttgagtcagacctgtaggcctgatagactgattaaaccacagaaggtgacctgctgagaaaagtggtacaaatactgggaaaaacctgctcttctgcgttaagtgggagacaatggcttcca

1 × 10^4^ cells HEK293T cells (n=6) were seeded in a 96-well plate and transfected at 70% confluency using lipofectamine 3000 with 200ng of either pGL4.73_5’UTR_tpx2; GQ mutant; GQ positive control or empty vector. 20 ng of empty firefly luciferase pGL4.13 plasmid was co-transfected as internal control. The following day, the cells were treated with either DMSO or CR-1-31-B (25nM) for 24 hours. The Dual-Luciferase® Reporter Assay (Promega) was performed according to the manufacturer’s instructions. The read-out was done on a GloMax®-Multi Detection System.

#### Patient-derived xenograft generation

Tumor tissue from two neuroblastoma cancer patients was collected under an approved IRB protocol (protocol #14-091). Tumor tissue was immediately minced, mixed (50:50) with matrigel (Corning, New York, NY) and implanted subcutaneously (s.c.) in the flanks of 6-8 weeks old female NOD/SCID/IL2Ry^-/-^ (NSG) mice (Jackson Laboratory, Bar Harbor, ME) to generate Patient Derived Xenografts (PDX) as previously described^27^. Mice were monitored daily, and models were transplanted in mice three times before being deemed established. PDX tumor histology was then confirmed by pathology review of H&E slides, and direct comparison to the corresponding patient slides.

#### In vivo treatment

Established PDXs were serially transplanted s.c. into the flank of 6-8 weeks old NSG mice as described above. Once tumors reached an average volume of 100-150 mm^3^, mice were randomized into treatment arms (n=5 mice per group) to receive either CR-1-31-B intraperitoneal (i.p.) at a dose of 0.25 mg/kg or vehicle control (5.2%Tween 80 and 2% DMSO) three times per week. Mice were observed daily throughout the treatment period for signs of morbidity and mortality. Tumors were measured twice weekly using calipers, and volume was calculated using the formula: length x width^2^ × 0.52. Body weight was also assessed twice weekly. After ∼4 weeks of treatment tumor samples were collected for further analysis. When the tumor size reached 1500 mm^3^, animals were euthanized and their tumors were harvested for further histology analysis. All *in vivo* studies were carried out under an approved IACUC protocol and in accordance with guidelines approved by the Memorial Sloan Kettering Cancer Center Research Animal Resource Center.

#### Shotgun proteomics

IMR-32 neuroblastoma cells were seeded in a 175cm² cell culture flask at a density of 7.5 × 10^6^ cells per flask. Cells were allowed to adhere for 48 hours, after which the medium was replaced by fresh medium and the treatment. For the CR-1-31-B condition, 3.334 nM (IC50, 72h) was added, while 0.1% of DMSO was added as the control condition. Cells were collected at 3h, 6h, 12h, 24h, 48h and 72h after treatment. Upon the selected harvesting time points, cells were scraped, centrifuged for 5 minutes at 1200 rpm, and the pellet was washed twice with ice-cold PBS.

Cell pellets were homogenized in 1 mL lysis buffer containing 8 M urea and 20 mM Hepes pH 8.0. Next, samples were sonicated with 3 pulses of 15 s at an amplitude of 20% using a 3 mm probe, with incubation on ice for 1 minute between pulses. After centrifugation for 15 minutes at 20,000 x g at room temperature (RT) to remove insoluble components, proteins were reduced by addition of 5 mM DTT and incubation for 30 minutes at 55°C and then alkylated by addition of 10 mM iodoacetamide and incubation for 15 minutes at RT in the dark. The protein concentration was measured using a Bradford assay (Bio-rad) and from each sample 100 µg protein was used to continue the protocol. Samples were diluted with 20 mM HEPES pH 8.0 to a urea concentration of 4 M and proteins were digested with 1 µg LysC (Wako) (1/100, w/w) for 4 hours at 37°C. Samples were further diluted to a urea concentration of 2 M and digested with 1 µg trypsin (Promega) (1/100, w/w) overnight at 37°C. The resulting peptide mixture was acidified by addition of 1% trifluoroacetic acid (TFA) and after 15 minutes incubation on ice, samples were centrifuged for 15 minutes at 1,780 x g at room temperature to remove insoluble components. Next, peptides were purified on OMIX C18 tips (Agilent). The tips were first washed 3 times with 200 µL pre-wash buffer (0.1% TFA in water/acetonitrile (ACN) (20:80, v/v)) and pre-equilibrated 5 times with 200 µL of solvent A (0.1% TFA in water/ACN (98:2, v/v)) before samples were loaded on the tip. After peptide binding, the tip was washed 3 times with 200 µL of solvent A and peptides were eluted twice with 150 µL elution buffer (0.1% TFA in water/ACN (40:60, v/v)). The combined elutions were transferred to HPLC inserts and dried in a vacuum concentrator. To verify if the samples from different groups were clustered together and to detect outlier samples PCA was performed using Python, on the LFQ intensity. Statistical testing was done using the 2-way ANOVA, in order to identify proteins that were significantly evolving through time and between condition. As to correct for multiple testing, a Benjamini-Hochberg correction was applied on the p-value of the 2-way ANOVA test. GSEA was performed on the genes ordered according to the differential expression statistic value (t). Signature scores were used for visualization of gene set enrichment. For gene ontology analysis, Enrichr^28^, was used with the default setting to identify enriched functional classes and biological processes of the identified proteins.

#### CUT&RUN

CUT&RUN coupled with high-throughput DNA sequencing was performed on isolated nuclei using Cutana pA/G-MNase (Epicypher, 15-1016) according to the manufacturer’s manual. Briefly, nuclei were isolated from 0.5M cells/sample in 100µL nuclear extraction buffer per sample and incubated with activated Concanavalin A beads for 10 min at 4°C while rotating. Nuclei were resuspended in 50µL antibody buffer containing a 1:100 dilution of each antibody (N-MYC and IgG) and kept in an elevated angle on a nutator at 4°C overnight. Next, targeted digestion and release was performed with 2.5 µL Cutana pA/GMNase (15-1116) and 100mM CaCl2 for 2 hours at 4°C on the nutator. After chromatin release by incubation on 37°C for 10 minutes, DNA was purified using the CUTANA DNA purification kit (14-0050) and eluted in 12µL of elution buffer. Sequencing libraries were prepared with the NEBNext Ultra II kit (Illumina, E7645), followed by paired-end sequencing on a Nextseq2000 using the NextSeq 2000 P2 Reagents 100 Cycles v3 (Illumina, 20046811). Prior to mapping to the human reference genome (GRCh37/hg19) with bowtie2 (v.2.3.1), quality of the raw sequencing data of CUT&RUN was evaluated using FastQC and adapter trimming was done using TrimGalore (v0.6.5). Quality of aligned reads were filtered using min MAPQ 30 and reads with known low sequencing confidence were removed using Encode Blacklist regions. For samples with a percentage duplicated reads higher then 10%, deduplication was performed using MarkDuplicates (Picard, v4.0.11). Peak calling was performed using MACS2 (v2.1.0) taking a q value of 0.05 as threshold and default parameters. Homer (v4.10.3) was used to perform motif enrichment analysis, with 200 bp around the peak summit as input. The R package igvR (v1.19.3) was used for visualization of the data upon RPKM normalization. All CUT&RUN data are available through the Gene Expression Omnibus (GEO) repository (GSE246722).

#### Ribosome profiling and RNA sequencing sample preparation

IMR-32 neuroblastoma cells were seeded in a 175cm² cell culture flask at a density of 7.5 × 10^6^ cells per flask. Cells were allowed to adhere for 48 hours, after which the medium was replaced by fresh medium and the treatment. For the CR-1-31-B condition, 3.334 nM (IC50) was added, 0.1% of DMSO was added for the control condition. After 6 hours of treatment, cycloheximide (CHX) was applied with a final concentration of 0.1 mg/mL prior to lysis. After 1 minute of incubation, the medium was aspirated and cells were rinsed with 10 mL of ice-cold PBS supplemented with CHX (0.1 mg/ml). Finally, cells were scraped in 5mL of PBS with CHX (0.1 mg/mL). The samples were centrifuged at 1000 rpm for 5 minutes, at 4°C.

#### RNA-sequencing drugging

RNA extraction was performed according to manufacturer’s instructions of the miRNeasy mini kit (Qiagen) including on-column DNase treatment. The concentration was measured by using Nanodrop (Thermo Scientific). RNA quality was determined with the Experion automated electrophoresis system (BioRad) prior to profiling. 250 ng of RNA isolated, as described above, was used as input for library preparation with the TruSeq small RNA library Kit (Illumina). During library preparation 15 PCR cycles were used. qPCR quantification of the libraries was done using the Kapa library quantification kit (Lightcycler 480 qPCR mix Kapa). RNA-Seq libraries were sequenced on the NextSeq 500 platform (Illumina) using the NextSeq 500 High Output kit V2.5 75 cycles single-end (Illumina). Sample and read quality were checked with FastQC (version 0.11.7). The FASTQ files were aligned to the *Homo sapiens* reference genome (GRCh38.96) and reads were counted using STAR (version 2.7.2b). Counts were normalized using edgeR (version 3.36.0) and differential expression analysis was performed using Limma-Voom (version 3.50.3).

#### Ribosome profiling

IMR-32 cells were grown in a 175cm² flask in complete RPMI 1640 medium at a cell density of 7.5 × 10^6^ and treated with CR-1-31-B (3.4 nM) or DMSO (0.1%). After 6 hours, cells were treated with CHX (0.1 mg/mL) before harvesting. Cell lysis of the cell pellets was performed using Mammalian Polysome buffer with CHX. The homogenized cell lysates were incubated for 10-30 minutes on ice before centrifugation (10 minutes, 18,000g, 4°C). The cleared supernatant was recovered. The lysates were quantified using the Qubit RNA High sensitivity kit (Fisher scientific). 15µg of the lysates were treated with 7.5 units RNase I, purified with the Microspin S-400 columns and the Zymo RNA clean and concentrator kit (Zymo research). The eluted samples were size selected (28-30 nt) on a 15% PAGE urea gels. Samples were 3’ dephosphorylated with T4 PNK followed by library prep using the Small RNA library prep kit (Lexogen). rRNA was blocked with LNA probes during the cDNA synthesis step of the library prep using the FastSelect H/M/R kit from Qiagen. Library amplification was done according to Lexogen small RNA kit. The PCR products were purified using Ampure XP beads. Library quality control was done on a DNA high sensitivity chip (BioAnalyzer Agilent, 5067-4626). Library concentrations were measured with qPCR according to Illumina’s protocol. Libraries were sequenced on 2 NextSeq500 high output runs with 76bp single reads.

Adaptor sequences (TGGAATTCTCGGGTGCCAAGGAACTCCAGTCAC) were trimmed from RiboSeq reads using the cutadapt 3.2 with parameters (-j 10 -m 20 --discard-untrimmed). rRNA-derived reads were identified and discarded by bowtie 2.3.4.1 with default settings. The gene reads count were computed by RSEM v1.2.28 with default settings except using STAR 2.7.1a aligner for mapping. The reference transcripts were generated using hg38 as reference genome and GENCODE v37 as gene annotation. Differential analysis was performed using the R package DESeq2 1.32.0. and genes were normalized using Transcripts Per Million (TPM) normalization.

To verify if the samples from different groups were clustered together and to detect outlier samples, Principal Component Analyses (PCAs) on rlog transformed counts were performed using the R statistical computing software. DESeq2 (1.32.0) was applied to RNA-Seq and RiboSeq counts in order to obtain the differential transcribed genes (DTG) and differential translation efficiency genes (DTEGs). The analysis was carried out as described in ^13^, in order to calculate translation efficiency, which is the number of ribosomes per gene normalized to transcripts abundance. GSEA was performed on the genes grouped by category, exclusive, forwarded, intensive, buffer and buffered special, were ordered according to the differential expression statistic value (t). Signature scores were used for visualization of gene set enrichment. For gene ontology analysis, Enrichr^28^, was used with the default setting to identify enriched functional classes.

#### MS analysis

The peptides were re-dissolved in 20 µL loading solvent A (0.1% TFA in water/ACN (98:2, v/v)) of which 4 µL was injected for LC-MS/MS analysis on an Ultimate 3000 RSLCnano system in-line connected to a Q Exactive HF mass spectrometer (Thermo). Trapping was performed at 10 μL/min for 4 min in loading solvent A on a 20 mm trapping column (made in-house, 100 μm internal diameter (I.D.), 5 μm beads, C18 Reprosil-HD, Dr. Maisch, Germany). The peptides were separated on an in-house produced column (75 µm x 500 mm), equipped with a laser pulled electrospray tip using a P-2000 Laser Based Micropipette Puller (Sutter Instruments), packed in-house with ReproSil-Pur basic 1.9 µm silica particles (Dr. Maisch). The column was kept at a constant temperature of 50°C. Peptides eluted using a non-linear gradient reaching 30% MS solvent B (0.1% FA in water/acetonitrile (2:8, v/v)) in 105 min, 55% MS solvent B in 145 min and 99% MS solvent B after 150 min at a constant flow rate of 250 nL/min. This was followed by a 10-minutes wash at 99% MS solvent B and re-equilibration with MS solvent A (0.1% FA in water). The mass spectrometer was operated in data-dependent mode, automatically switching between MS and MS/MS acquisition for the 16 most abundant ion peaks per MS spectrum. Full-scan MS spectra (375-1500 m/z) were acquired at a resolution of 60,000 in the Orbitrap analyzer after accumulation to a target value of 3,000,000. The 16 most intense ions above a threshold value of 13,000 were isolated (isolation window of 1.5 m/z) for fragmentation at a normalized collision energy of 28% after filling the trap at a target value of 100,000 for maximum 80 ms. MS/MS spectra (145-2,000 m/z) were acquired at a resolution of 15,000 in the Orbitrap analyzer. Data analysis was performed with MaxQuant (version 1.6.1.0) using the Andromeda search engine with default search settings including a false discovery rate set at 1% on both the peptide and protein level. Spectra were searched against the human proteins in the Uniprot/Swiss-Prot database (database release version of April 2018 containing 20,243 human protein sequences, downloaded from www.uniprot.org). The mass tolerance for precursor and fragment ions was set to 4.5 and 20 ppm, respectively, during the main search. Enzyme specificity was set as C-terminal to arginine and lysine, also allowing cleavage at proline bonds with a maximum of two missed cleavages. Variable modifications were set to oxidation of methionine residues and acetylation of protein N-termini, while carbamidomethylation of cysteine residues was set as fixed modification. Matching between runs was enabled with a matching time window of 1 minute and an alignment time window of 20 minutes. Only proteins with at least one unique or razor peptide were retained leading to the identification of 5,671 proteins. Proteins were quantified by the MaxLFQ algorithm integrated in the MaxQuant software. A minimum ratio count of two unique or razor peptides was required for quantification. Further data analysis was performed with the Perseus software (version 1.5.5.3) after loading the proteingroups file from MaxQuant. Proteins only identified by site, reverse database hits and possible contaminants were removed and replicate samples were grouped. Proteins with less than three valid values in at least one group were removed and missing values were imputed from a normal distribution around the detection limit, leading to a list of 4,452 quantified proteins that was used for further data analysis. Then, t-tests were performed (FDR=0.05 and s0=1) to compare samples and generate the volcano plots. Additionally, to reveal proteins of which the expression level was significantly affected between the different conditions, sample groups were defined based on the time (3h, 6h, 12h, 24h, 48h and 72h) and treatment (control vs. Treated) and a two-way ANOVA test was performed to compare the intensities of the proteins in the time group with the treatment group. For each protein, this test calculated a p-value (actually -log p-value) for time, a p-value for treatment and a p-value for the interaction of time and treatment. 1,555 proteins with a p-value <0.05 in at least one of these three groups were considered to be significantly regulated. The intensities of these proteins were plotted in two heat maps after non-supervised hierarchical clustering.

#### GQ analyses

Transcript IDs were essential for identifying the correct 5’UTR of each expressed gene. To achieve this, kallisto quant (version 0.48.0) was utilized for mapping the transcript IDs, employing hg38 index files. Subsequently, the Ensemble gene names were matched with the transcript IDs via the R package biomaRt (version 2.58.2). Genes with the highest transcript counts were retained and 5’UTR information was added via biomaRt. These expressed transcripts were then integrated with the different classes: forwarded, buffered special, exclusive, and the background translation efficiency list.

Potential GQ DNA motifs analysis in the 5’UTR region of the genes in the different classes (forwarded, buffered special and exclusive) was performed using the tool ‘Quadparser’ using an adapted regular expression “([gG]{3,}\w{1,10}){3,}[gG]{3,]” as described in ^14^. Initially, the 5’UTRs were converted from a bed format to a fasta format using BEDTools (version 2.31.0) getfasta function.

Motif enrichment itself was performed using ‘Simple Enrichment Analysis’ (SEA) hosted by The MEME Suite^29^ using the HOCOMOCO Human (version 11 FULL) database. The sequence logo was created using seqLog^30^ (version 1.66.0) library in R.

All scripts used in these analyses are available on GitHub: https://github.com/PPOLLabGhent/CR-1-31-B_TranslationInsights

#### siRNA mediated TPX2 knockdown

IMR-32 cells were transfected using the Neon Transfection Kit (ThermoFisher) with siRNAs for TPX2 (Ambion, s22745 (siTPX2 #45), s22746 (siTPX2 #46) and s22747 (siTPX2 #47)) or scrambled siRNA (Horizon Discovery Limited, D-001810-10-05). After transfection, cells were seeded in 25 cm^2^ culture flasks at a density of 2 × 10^6^ cells (for protein and RNA isolation), in a 96-well plate at a density of 3 × 10^4^ cells (for apoptosis assays) and in a 384-well plate at a density of 3 × 10^3^ cells. The 384-well plates were used for drugging with prexasertib or CR-1-31-B with a concentration of 2nM. Cell confluency was monitored by IncuCyte Live-Cell Imaging. Each image was analyzed through the IncuCyte Software and cell proliferation was determined by the area (percentage of confluence) of cell images over time. 48h after transfection, apoptosis levels were measured from the 96-wells plate using the Caspase-Glo 3/7 Assay (Promega). The protocol was adapted to add 50 µL of caspase reagent per well. The average luminescence was normalized to the cell-occupied area, followed by normalization to the scrambled siRNA control wells. Protein levels were measured using the method described in the Western Blot analysis section. RNA was isolated using the protocol from the miRNeasy mini kit (Qiagen), including on-column DNase treatment. The RNA concentration was measured using the NanoDrop (Isogen Lifescience). Complementary DNA (cDNA) was produced using the iScript Advanced cDNA Synthesis kit (Bio-Rad). After cDNA production, the PCR mastermix was made containing 5 ng cDNA, 2.5 µL SsoAdvanced Universal SYBR Green Supermix (Bio-Rad) and 0.25 µL forward and reverse primer. *Forward primer TPX2*: *TGTTGTGGGTGTTCCTGAAA, Reverse primer TPX2: CGAGAAAGGGCATATTTCCA* The RT-qPCR was performed using the LC-480 (Roche) and the gene expression analysis was performed using Qbase+. The average gene expression of *TPX2* was normalized to the scrambled siRNA control.

For all, statistics were performed using the GraphPad Prism software (version 10.0.1), the tests used were one-way ANOVA with Dunnet’s multiple comparison test. Statistical significance was determined by a p-value of ≤0.05.

#### Western blot analysis

Cells were seeded in 75 cm² cell culture flasks at a density of 2.5 × 10^6^ cells and allowed to adhere for 48 hours. After 48 hours, the medium was refreshed with medium containing at 3.334 nM of CR-1-31-B (IC50) or 0.1% DMSO. Cells were harvested by scraping in ice-cold PBS, followed by centrifugation at 2500 rpm for 5 minutes at 4°C. The pellet was rinsed, once with ice-cold PBS. Cells were lysed in cold RIPA buffer (250mg sodium deoxycholate, 150mM NaCl, 50mM Tris-HCl pH 7.5, 0.01% SDS solution, 0.1% NP-40) supplemented with protease and phosphatase inhibitors (Roche). Samples were rotated for 1 hour at 4°C.The cleared lysates were collected and centrifuged at 10,000 rpm for 10 minutes at 4°C. Protein concentrations were determined using the protocol of the Pierce^TM^ BCA Protein Assay Kit (ThermoFisher). The lysates were denaturized prior to loading on a gel, through 5 times Laemmli denaturation buffer supplemented with β-mercaptoethanol (25mL 10% SDS solution, 21.25mL 100% glycerol, 7.75mL 1M Tris-HCl pH 6.8, 0.0125mg bromophenol blue). 30µg of protein extracts were loaded on 10% SDS-PAGE gels with 10x Tris/glycine/SDS buffer and run for 1 hour at 130 V. Samples were blotted on nitrocellulose or PVDF membranes in 10% of 10x Tris/Glycine buffer and 20% of methanol. The membranes were blocked during 1 hour in 5% milk or 5% BSA in TBS-T. Primary antibody incubations were done in blocking buffer overnight at 4°C. Blots were washed 3 times with TBST before the incubation for 1 hour of secondary antibodies. Before visualization, the blots were washed again 3 times with TBST. The immunoblots were visualized by using the enhanced chemiluminescent West Femto or West Dura (Bio-Rad). The protein quantification analysis of the generated blots was performed through ImageJ software, where the area from each protein was normalized to the loading protein in respect to each blot. The antibodies used were cleaved PARP (1:2000), LH2AX (1:2000), vinculin (1:10,000) and TPX2 (1:10,000), anti-rabbit IgG (1:5000) and anti-mouse IgG (1:5000).

#### Combination treatment experiments

IMR-32 cells were seeded in 384-well plates at a density of 2000 cells/well and allowed to adhere overnight. Next, cells were treated with Prexasertib or CR-1-31-B alone or in a combination matrix using the Tecan D300e Digital Dispenser. Cell confluency was monitored by IncuCyte Live-Cell Imaging. Each image was analyzed through the IncuCyte Software and cell proliferation was determined by the area (percentage of confluence) of cell images over time. Synergism was determined via the algorithm HTSplotter^31^, using the Bliss Index (BI) score. Graphs were made using GraphPad (version 10.0.1).

### Data availability statement

All data generated during this study were deposited on NCBI GEO database, the RNA-sequencing and ribosome sequencing of the CR-3-1-B drugging (GSE216193), RNA-sequencing of siTPX2 (GSE246725) and CUT&RUN (GSE246722). The mass spectrometry proteomics data have been deposited to the ProteomeXchange Consortium via the PRIDE^32^ partner repository with the dataset identifier PXD044871. Original western blot images and STR genotype files were deposited to Mendeley (DOI: 10.17632/8btwfg6hm9.1).

Modelling computer scripts are available on GitHub: https://github.com/PPOLLabGhent/CR-1-31-B_TranslationInsights

## References

1. Pelletier, J., Thomas, G., and Volarević, S. (2018). Ribosome biogenesis in cancer: new players and therapeutic avenues. Nat Rev Cancer 18, 51–63.

2. Bhat, M., Robichaud, N., Hulea, L., Sonenberg, N., Pelletier, J., and Topisirovic, I. (2015). Targeting the translation machinery in cancer. Preprint at Nature Publishing Group, 10.1038/nrd4505 10.1038/nrd4505.

3. Wolfe, A.L., Singh, K., Zhong, Y., Drewe, P., Rajasekhar, V.K., Sanghvi, V.R., Mavrakis, K.J., Jiang, M., Roderick, J.E., Van der Meulen, J., et al. (2014). RNA G-quadruplexes cause eIF4A-dependent oncogene translation in cancer. Nature 513, 65–70. 10.1038/nature13485.

4. Singh, K., Lin, J., Lecomte, N., Mohan, P., Gokce, A., Sanghvi, V.R., Jiang, M., Grbovic-Huezo, O., Burčul, A., Stark, S.G., et al. (2021). Targeting eIF4A-dependent translation of KRAS signaling molecules. Cancer Res 81, 2002–2014. 10.1158/0008-5472.CAN-20-2929.

5. Chan, K., Robert, F., Oertlin, C., Kapeller-Libermann, D., Avizonis, D., Gutierrez, J., Handly-Santana, A., Doubrovin, M., Park, J., Schoepfer, C., et al. (2019). eIF4A supports an oncogenic translation program in pancreatic ductal adenocarcinoma. Nat Commun 10. 10.1038/s41467-019-13086-5.

6. Skofler, C., Kleinegger, F., Krassnig, S., Birkl-Toeglhofer, A.M., Singer, G., Till, H., Benesch, M., Cencic, R., Porco, J.A., Pelletier, J., et al. (2021). Eukaryotic translation initiation factor 4AI: A potential novel target in neuroblastoma. Cells 10, 1–16. 10.3390/cells10020301.

7. Xue, C., Gu, X., Li, G., Bao, Z., and Li, L. (2021). Expression and Functional Roles of Eukaryotic Initiation Factor 4A Family Proteins in Human Cancers. Preprint at Frontiers Media S.A., 10.3389/fcell.2021.711965 10.3389/fcell.2021.711965.

8. Shen, L., Pugsley, L., Cencic, R., Wang, H.C., Robert, F., Naineni, S.K., Sahni, A., Morin, G., Zhang, W., Nijnik, A., et al. (2021). A forward genetic screen identifies modifiers of rocaglate responsiveness. Scientific Reports 2021 11:1 11, 1–13. 10.1038/s41598-021-97765-8.

9. Gupta, S. V., Sass, E.J., Davis, M.E., Edwards, R.B., Lozanski, G., Heerema, N.A., Lehman, A., Zhang, X., Jarjoura, D., Byrd, J.C., et al. (2011). Resistance to the Translation Initiation Inhibitor Silvestrol is Mediated by ABCB1/P-Glycoprotein Overexpression in Acute Lymphoblastic Leukemia Cells. AAPS J 13, 357–364. 10.1208/S12248-011-9276-7.

10. Decaesteker, B., Denecker, G., Van Neste, C., Dolman, E.M., Van Loocke, W., Gartlgruber, M., Nunes, C., De Vloed, F., Depuydt, P., Verboom, K., et al. (2018). TBX2 is a neuroblastoma core regulatory circuitry component enhancing MYCN/FOXM1 reactivation of DREAM targets. Nat Commun 9. 10.1038/s41467-018-06699-9.

11. Vanhauwaert, S., Decaesteker, B., De Brouwer, S., Leonelli, C., Durinck, K., Mestdagh, P., Vandesompele, J., Sermon, K., Denecker, G., Van Neste, C., et al. (2018). In silico discovery of a FOXM1 driven embryonal signaling pathway in therapy resistant neuroblastoma tumors. Sci Rep 8. 10.1038/s41598-018-35868-5.

12. Grant, G.D., Brooks, L., Zhang, X., Mahoneya, J.M., Martyanov, V., Wood, T.A., Sherlock, G., Cheng, C., and Whitfield, M.L. (2013). Identification of cell cycle-regulated genes periodically expressed in U2OS cells and their regulation by FOXM1 and E2F transcription factors. Mol Biol Cell 24, 3634–3650. 10.1091/mbc.E13-05-0264.

13. Chothani, S., Adami, E., Ouyang, J.F., Viswanathan, S., Hubner, N., Cook, S.A., Schafer, S., and Rackham, O.J.L. (2019). deltaTE: Detection of Translationally Regulated Genes by Integrative Analysis of Ribo-seq and RNA-seq Data. Curr Protoc Mol Biol 129. 10.1002/cpmb.108.

14. Puig Lombardi, E., and Londoño-Vallejo, A. (2020). A guide to computational methods for G-quadruplex prediction. Nucleic Acids Res 48, 1. 10.1093/NAR/GKZ1097.

15. Molenaar, J.J., Van Sluis, P., Boon, K., Versteeg, R., and Caron, H.N. (2003). Rearrangements and increased expression of cyclin D1 (CCND1) in neuroblastoma. Genes Chromosomes Cancer 36, 242–249. 10.1002/gcc.10166.

16. Decaesteker, B., Louwagie, A., Loontiens, S., De Vloed, F., Bekaert, S.L., Roels, J., Vanhauwaert, S., De Brouwer, S., Sanders, E., Berezovskaya, A., et al. (2023). SOX11 regulates SWI/SNF complex components as member of the adrenergic neuroblastoma core regulatory circuitry. Nat Commun 14. 10.1038/s41467-023-36735-2.

17. Molenaar, J.J., Koster, J., Zwijnenburg, D.A., Van Sluis, P., Valentijn, L.J., Van Der Ploeg, I., Hamdi, M., Van Nes, J., Westerman, B.A., Van Arkel, J., et al. (2012). Sequencing of neuroblastoma identifies chromothripsis and defects in neuritogenesis genes. Nature 483, 589–593. 10.1038/NATURE10910.

18. Ognibene, M., Podestà, M., Garaventa, A., and Pezzolo, A. (2019). Role of GOLPH3 and TPX2 in neuroblastoma DNA damage response and cell resistance to chemotherapy. Int J Mol Sci 20. 10.3390/ijms20194764.

19. De Wyn, J., Zimmerman, M.W., Weichert-leahey, N., Nunes, C., Cheung, B.B., Abraham, B.J., Beckers, A., Volders, P.J., Decaesteker, B., Carter, D.R., et al. (2021). Meis2 is an adrenergic core regulatory transcription factor involved in early initiation of th-mycn-driven neuroblastoma formation. Cancers (Basel) 13. 10.3390/cancers13194783.

20. Koppen, A., Ait-Aissa, R., Hopman, S., Koster, J., Haneveld, F., Versteeg, R., and Valentijn, L.J. (2007). Dickkopf-1 is down-regulated by MYCN and inhibits neuroblastoma cell proliferation. Cancer Lett 256, 218–228. 10.1016/J.CANLET.2007.06.011.

21. Cantor, S. (2019). TPX2 joins 53BP1 to maintain DNA repair and fork stability. Preprint at Rockefeller University Press, 10.1083/jcb.201812142 10.1083/jcb.201812142.

22. Byrum, A.K., Carvajal-Maldonado, D., Mudge, M.C., Valle-Garcia, D., Majid, M.C., Patel, R., Sowa, M.E., Gygi, S.P., Wade Harper, J., Shi, Y., et al. (2019). Mitotic regulators TPX2 and Aurora A protect DNA forks during replication stress by counteracting 53BP1 function. Journal of Cell Biology 218, 422–432. 10.1083/jcb.201803003.

23. Neumann, J., Boerries, M., Köhler, R., Giaisi, M., Krammer, P.H., Busch, H., and Li-Weber, M. (2014). The natural anticancer compound rocaglamide selectively inhibits the G1-S-phase transition in cancer cells through the ATM/ATR-mediated Chk1/2 cell cycle checkpoints. Int J Cancer 134, 1991–2002. 10.1002/ijc.28521.

24. King, D., Li, X.D., Almeida, G.S., Kwok, C., Gravells, P., Harrison, D., Burke, S., Hallsworth, A., Jamin, Y., George, S., et al. (2020). MYCN expression induces replication stress and sensitivity to PARP inhibition in neuroblastoma. Oncotarget 11, 2141–2159. 10.18632/ONCOTARGET.27329.

25. Volegova, M.P., Brown, L.E., Banerjee, U., Dries, R., Sharma, B., Kennedy, A., Porco, J.A., and George, R.E. (2024). The MYCN 5′ UTR as a therapeutic target in neuroblastoma. bioRxiv, 2024.02.20.581230. 10.1101/2024.02.20.581230.

26. Stockwell, S.R., Scott, D.E., Fischer, G., Guarino, E., Rooney, T.P.C., Feng, T.-S., Moschetti, T., Srinivasan, R., Alza, E., Asteian, A., et al. (2023). Selective inhibitors of the Aurora A-TPX2 protein-protein interaction exhibit in vivo efficacy as targeted anti-mitotic agents. bioRxiv, 2023.03.22.533679. 10.1101/2023.03.22.533679.

27. Mattar, M., McCarthy, C.R., Kulick, A.R., Qeriqi, B., Guzman, S., and de Stanchina, E. (2018). Establishing and maintaining an extensive library of patient-derived xenograft models. Front Oncol 8. 10.3389/fonc.2018.00019.

28. Xie, Z., Bailey, A., Kuleshov, M. V, Clarke, D.J.B., Evangelista, J.E., Jenkins, S.L., Lachmann, A., Wojciechowicz, M.L., Kropiwnicki, E., Jagodnik, K.M., et al. (2021). Gene Set Knowledge Discovery with Enrichr. Curr Protoc 1, e90. 10.1002/cpz1.90.

29. Bailey, T.L., and Grant, C.E. (2018). SEA: Simple Enrichment Analysis of motifs. 10.1101/2021.08.23.457422.

30. Bembom, O., and Ivanek, R. (2023). Bioconductor - seqLogo. 10.18129/B9.bioc.seqLogo.

31. Nunes, C., Anckaert, J., De Vloed, F., De Wyn, J., Durinck, K., Vandesompele, J., Speleman, F., and Vermeirssen, V. (2024). HTSplotter: An end-to-end data processing, analysis and visualisation tool for chemical and genetic in vitro perturbation screening. PLoS One 19. 10.1371/JOURNAL.PONE.0296322.

32. Perez-Riverol, Y., Bai, J., Bandla, C., García-Seisdedos, D., Hewapathirana, S., Kamatchinathan, S., Kundu, D.J., Prakash, A., Frericks-Zipper, A., Eisenacher, M., et al. (2022). The PRIDE database resources in 2022: a hub for mass spectrometry-based proteomics evidences. Nucleic Acids Res 50, D543–D552. 10.1093/NAR/GKAB1038.

