## Supplementary figures and images for "5’UTR translational inhibition of neuroblastoma dependency factors using the CR-1-31-B rocaglate"

### Supplementary Figure 1

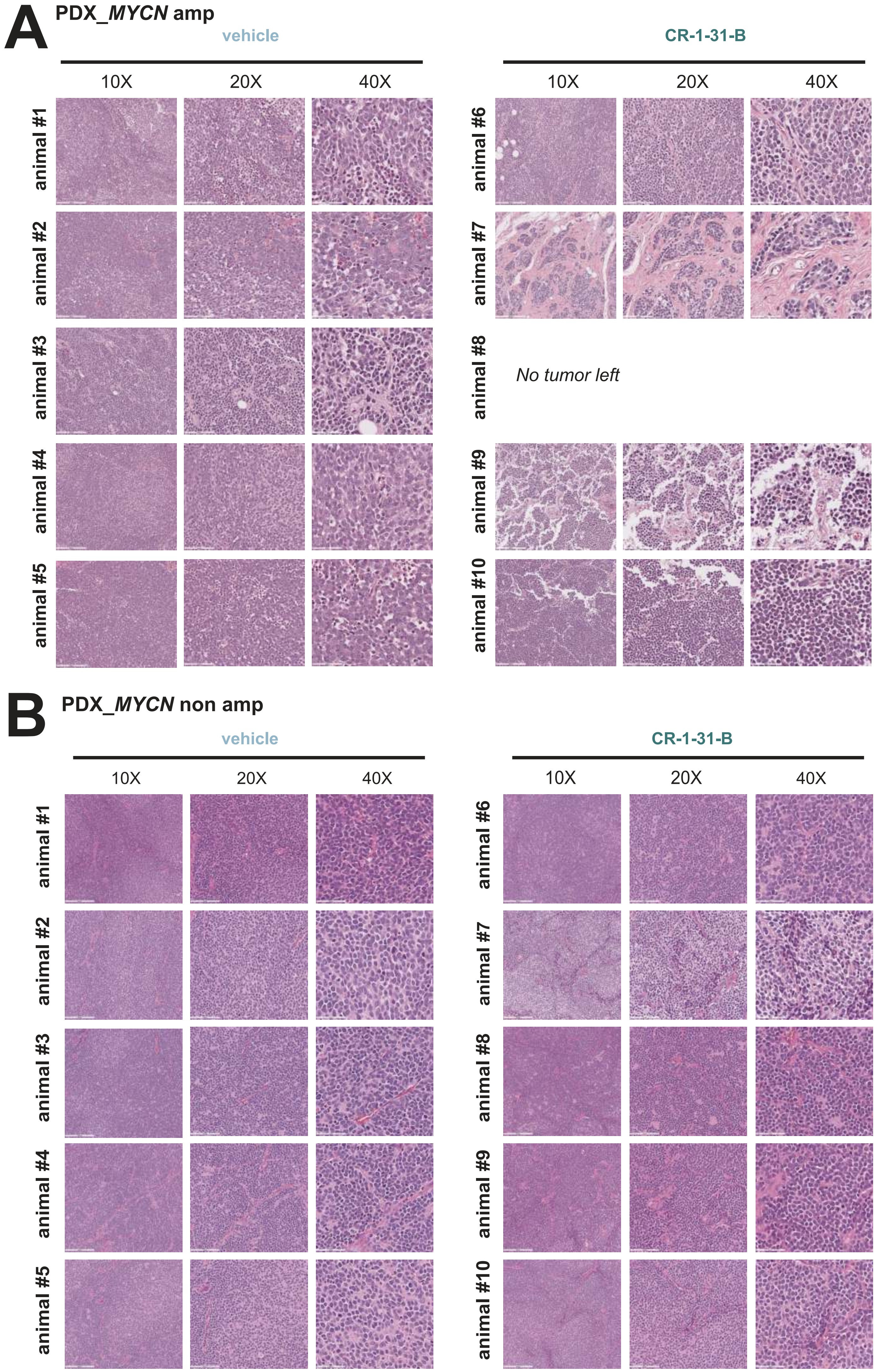

### Supplementary Figure 2

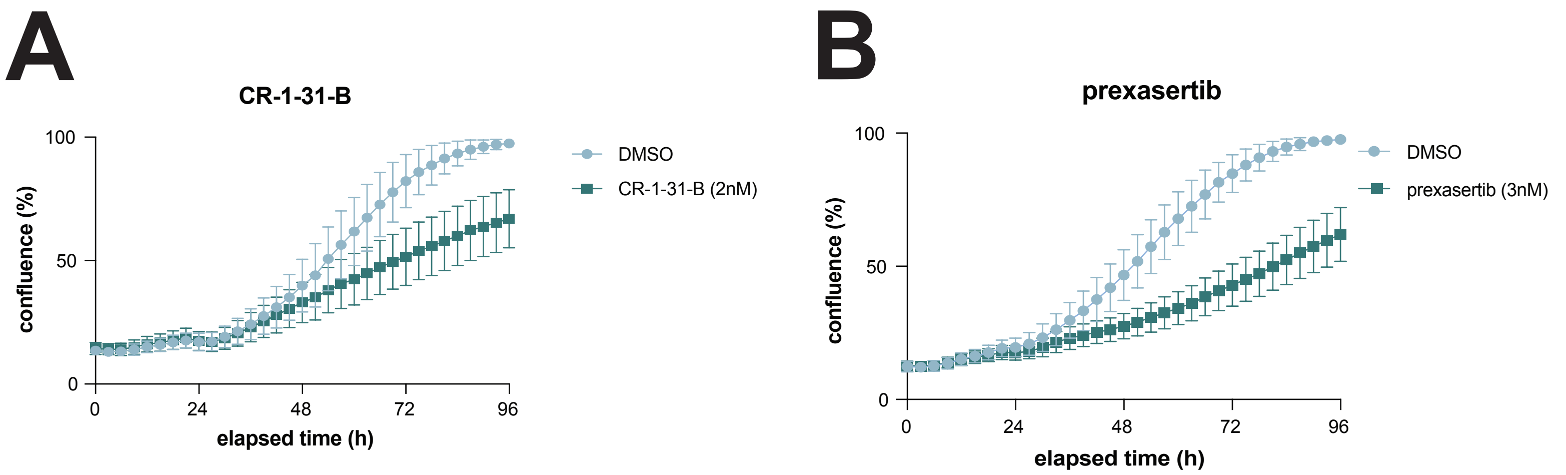
